# Progressive muscle metabolic reprogramming in asymptomatic ALS gene mutation carriers

**DOI:** 10.64898/2025.12.15.694300

**Authors:** Céline Buon, Dimitrije Milunov, Delphine Sapaly, Timothée Lenglet, Maria del Mar Amador, Elsie Piller, Sabrina Bendris, Flore Cheguillaume, Rasha Slika, Eleni Siopi, Stéphanie Bauché, Isabelle Le Ber, François Salachas, Foudil Lamari, Anthony Behin, Teresinha Evangelista, Baptiste Périou, François-Jérôme Authier, Bertrand Fontaine, Soham Saha, Frédéric Charbonnier, Laure Weill, Gaëlle Bruneteau

## Abstract

Amyotrophic lateral sclerosis (ALS) is a rapidly fatal neurodegenerative disorder characterized by motor neuron loss leading to extensive paralysis. There is emerging evidence that the disease involves a prolonged presymptomatic period during which motor function is preserved. Understanding the molecular mechanisms involved is key as the presymptomatic phase represents a critical window of opportunity for early intervention. Using RNA sequencing, we investigated changes in gene expression patterns in the skeletal muscle of ten asymptomatic carriers of ALS mutations (8 C9ORF72 expansion carriers and 2 SOD1 mutation carriers). We found that specific modifications of gene expression profiles are present in skeletal muscle before asymptomatic ALS gene carriers exhibit biomarker changes predictive of phenoconversion. We identified insulin signaling, AMPK signaling, and thermogenesis pathways, together with the TCA cycle as the main contributors to the dysregulated muscle transcriptome. Our data suggest that this metabolic reprogramming of skeletal muscle develops progressively during the transition to phenoconversion, characterized by a gradual increase in the expression of *SREBF1* which encodes SREPB1, the key transcriptional regulator of lipid synthesis, in parallel with the progressive activation of AMPK and insulin signaling pathways. Our findings are consistent with a progressive enhancement in fatty acid metabolism and oxidative capacity in skeletal muscle, followed by a decline in oxidative phosphorylation efficiency as phenoconversion approaches. Evidence of muscle metabolic reprogramming in ALS long before motor onset identifies the dysregulation of muscle energy homeostasis as a critical early event in ALS pathogenesis.

**One sentence summary:** Skeletal muscle from individuals at elevated genetic risk for ALS/FTD undergoes progressive metabolic reprogramming far before disease onset.

## INTRODUCTION

Amyotrophic lateral sclerosis (ALS) is a progressive neurodegenerative disorder involving motor neurons, resulting in rapidly progressive muscle paralysis and death from respiratory failure usually in 3-5 years. ALS is both clinically and genetically heterogeneous with a variety of motor phenotypes (*1*), about 10% of cases being familial and 90% sporadic.

ALS may have a prolonged presymptomatic period during which compensatory biological mechanisms maintain motor function (*2*). In asymptomatic ALS mutation carriers, levels of neurofilament fragments - including neurofilament light chain (NfL) - are known to increase in serum and cerebrospinal fluid from several months up to 5 years prior to phenoconversion to ALS (*3, 4*).

Beyond the nervous system, ALS is also a systemic disorder, including dysregulation of energy homeostasis, which strongly influences prognosis (*5*). This involves an increase in whole-body energy expenditure (*6, 7*) and alterations in skeletal muscle metabolism with greater reliance on fat oxidation (*8, 9*). Importantly, several indicators point to early, presymptomatic alterations of energy metabolism in ALS (*10*). Specific carbohydrate and lipid profiles have been reported to be associated with an increase or decrease in risk of developing the disease (*11, 12*). Changes in body composition, including loss of metabolically active cells, have also been reported in asymptomatic ALS mutation carriers at a stage when NfL levels remain within the normal range (*13*). The triggers for the switch between hypo- and hypermetabolic states remain to be determined. Furthermore, dysregulation of muscle cholesterol homeostasis has been reported in asymptomatic ALS mutation carriers, potentially contributing to the metabolic defect characteristic of ALS muscle (*14*).

In the present study, we used RNA sequencing to investigate muscle gene expression patterns in ten asymptomatic carriers of ALS gene mutations (8 C9ORF72 expansion carriers and 2 SOD1 mutation carriers). We found that specific modifications of muscle gene expression profiles can be detected before ALS gene carriers exhibit changes in classic biomarkers of neurodegeneration (i.e. increase in blood NfL levels, motor neuron loss assessed by Motor Unit Number Index, MUNIX). By combining advanced bioinformatics approaches, we identified the most influential gene clusters and built a pathophysiological model in which asymptomatic ALS gene carriers were stratified according to their muscle gene expression profiles.

## RESULTS

### Demographic characteristics and neurophysiological findings

Ten asymptomatic carriers of an ALS-causing gene mutation in C9ORF72 (n=8) or SOD1 (n=2) were enrolled in the PRE-ALS study (mean age 48,4 +/-11.4 [31-67], 7 females/3 males, Table 1, Table S1). Baseline characteristics of study participants have been described in part elsewhere (*14*). At enrollment, there was no detectable difference in deltoid CMAP and MUNIX in asymptomatic ALS mutation carriers compared with age- and sex-matched controls (Table S1). The CMAP and MUNIX sum-scores were also comparable at study entry between asymptomatic ALS mutation carriers and controls. There was no significant change in CMAP and MUNIX scores between baseline and the 18-month follow-up visit (Table S1). At the end of study follow-up, none of the PRE-ALS at-risk individuals presented clinical signs or symptoms indicative of phenoconversion.

**Table 1.**
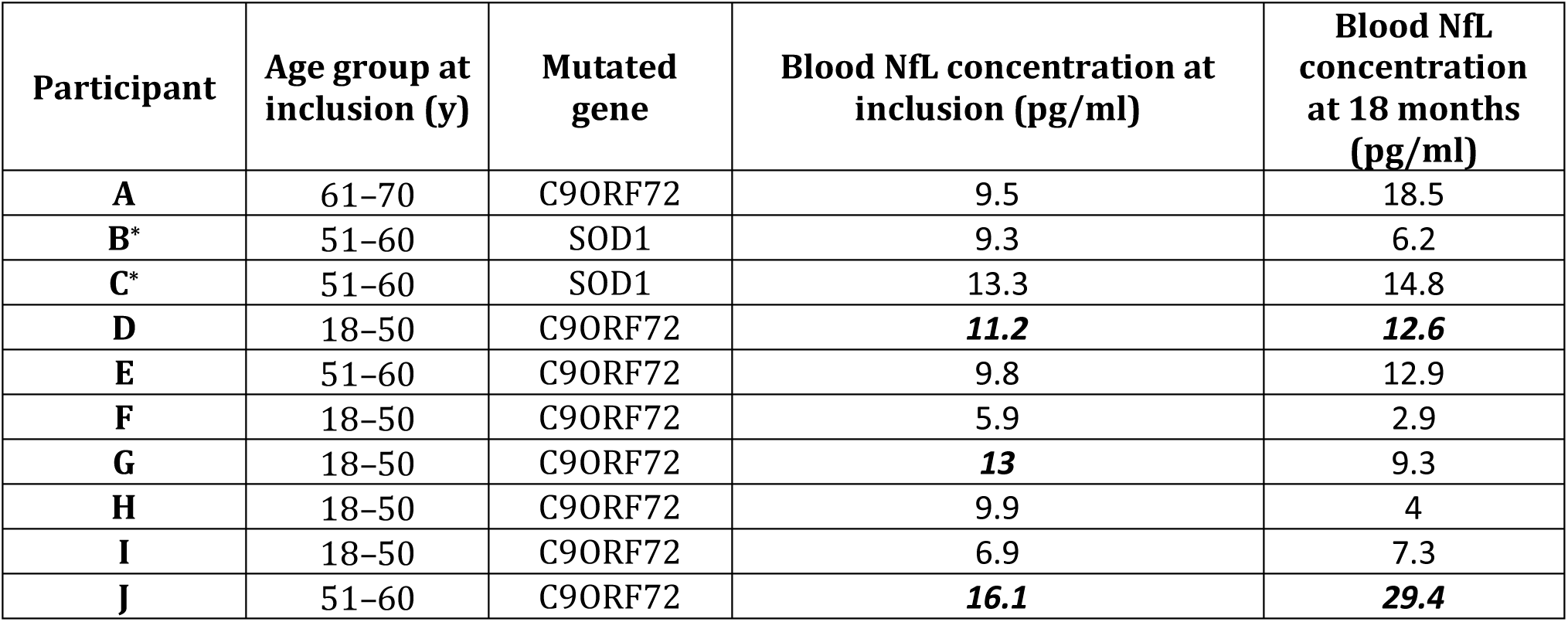
Nfl Levels in the ALS mutation carrier individuals. The following cut-off limits were used based on reference values previously reported in healthy individuals (*15*): 10 pg/mL in the group between ages of 18–50, 15 pg/mL in the ages ranging from 51–60, and 20 pg/mL in the group ranging between 61–70. Values in bold italics are above normal values for age. *Participants B and C were unaffected members within the same family carrying a c.418A>G (p.N139D) SOD1 mutation

| Participant | Age group at inclusion (y) | Mutated gene | Blood NfL concentration at inclusion (pg/ml) | Blood NfL concentration at 18 months (pg/ml) |
| --- | --- | --- | --- | --- |
| **A** | 61–70 | C9ORF72 | 9.5 | 18.5 |
| **B\*** | 51–60 | SOD1 | 9.3 | 6.2 |
| **C\*** | 51–60 | SOD1 | 13.3 | 14.8 |
| **D** | 18–50 | C9ORF72 | **11.2** | **12.6** |
| **E** | 51–60 | C9ORF72 | 9.8 | 12.9 |
| **F** | 18–50 | C9ORF72 | 5.9 | 2.9 |
| **G** | 18–50 | C9ORF72 | **13** | 9.3 |
| **H** | 18–50 | C9ORF72 | 9.9 | 4 |
| **I** | 18–50 | C9ORF72 | 6.9 | 7.3 |
| **J** | 51–60 | C9ORF72 | **16.1** | **29.4** |

### Neurofilament light chain concentrations

At inclusion (i.e. at the time of the muscle biopsy), three participants (D, G and J) had blood NfL values slightly above the cut-off value for their respective age group (*15*) (Table 1). For participant D, blood NfL concentration was just above the upper limit of normal at baseline and at the end of follow-up, with a 12.5% increase between the two measurements. For participant G, blood NfL levels had returned to the normal range at 18 months. The NfL level of participant J increased during follow-up, with an increment of more than 80% after 18 months compared to baseline.

### Transcriptomic analysis identifies early muscle alterations in asymptomatic ALS gene carriers

We performed RNA sequencing to analyze gene expression in the deltoid of PRE-ALS participants compared to deltoid samples from 10 age- and sex-matched control subjects (8 females / 2 males, mean age 51.4+/-7.2 [44-62], Table S2). Comparing the PRE-ALS group to controls, we observed greater variability and orthogonal expression patterns in gene expression (Fig. 1A, B). Principal component analysis (PCA) revealed that the first two components (PC1 and PC2) accounted for 32% and 11% of the total variance, respectively (Fig. 1A). Volcano plots (Fig. 1C) identified 659 differentially expressed genes (DEGs) at a stringent threshold of p<0.001 and |logFC| > 0.5. For ontology analysis, we relaxed the p-value threshold to p<0.05 to capture the collective impact of genes on cellular function, yielding 910 DEGs. These DEGs were involved in broad biological functions like metabolic processes, cellular processes, and cellular localizations (Fig. 1D). Specifically, upregulated genes (n = 434) were enriched in pathways such as GTP hydrolysis, muscle structure development, muscle contraction and mitochondrial processes (Fig. 1E), while downregulated genes (n = 476) were implicated in processes such as cellular respiration, actin filament processes, glycolysis, and proton membrane transport (Fig. 1F).

**Figure 1.**
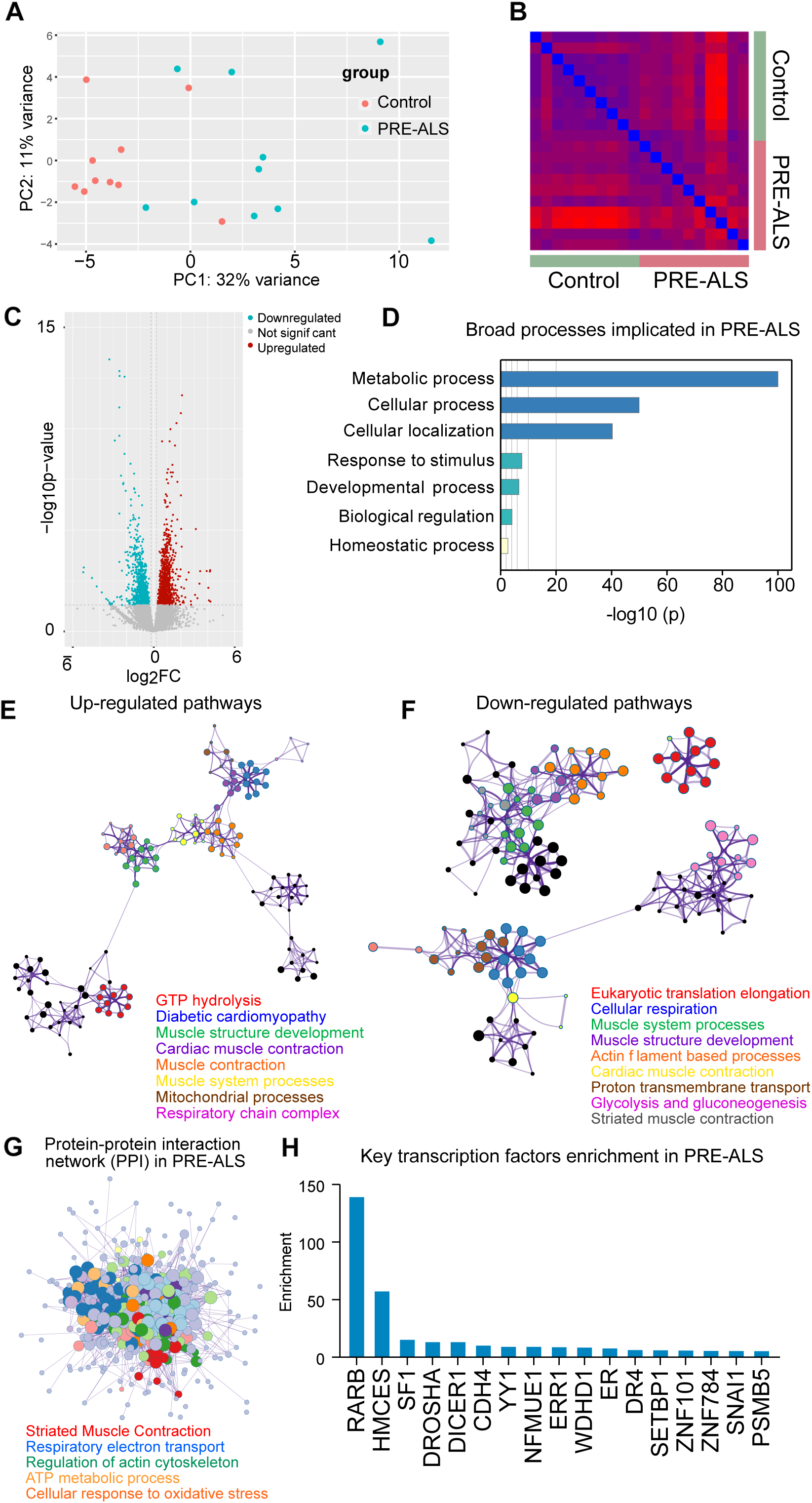
RNA-Seq analysis from controls and pre-ALS participants muscles shows differentially expressed genes. (A) Principal component analysis (PCA) along the first two principal components (PC1: 32% variance; PC2: 11% variance) (B) Correlation matrix from the differential expression patterns in PRE-ALS participants compared to controls. (C) Volcano plots showing the DEG (the log(FC) threshold is kept at ±0.5 (upregulation in red and downregulation in blue, p-value adjusted threshold is at p < 0.001; gray, non-significant transcripts). (D) Bar plot of enriched molecular processes identified through gene ontology analysis of upregulated genes in the PRE-ALS cohort compared to controls. (E) Gene network representation generated from key upregulated genes in the pre-ALS cohort compared to controls, highlighting muscle-specific pathways. (F) Gene network representation generated from key downregulated genes in the PRE-ALS cohort compared to controls, highlighting muscle-specific pathways. (G) A protein-protein interaction graph generated from ontology analysis of the upregulated genes in the PRE-ALS cohort compared to controls. (H) Identification and enrichment of key transcription factors in PRE-ALS cohort from the protein-protein interaction functional module analysis.

Protein-protein interaction enrichment analysis of the upregulated genes revealed functionally significant clusters, with key associated pathways including muscle contraction, respiratory electron transport, regulation of actin cytoskeleton, ATP metabolic process, cellular response to oxidative stress (Fig. 1G). Further network analysis using Metascape (*16*) identified several enriched transcription factors, notably RARB, HMCES, SF1, and miRNA processors DROSHA and DICER1 (Fig. 1H). Retinoic acid receptor beta (RARB), a member of the nuclear receptor superfamily known to mediate vitamin A-dependent transcriptional regulation of cellular differentiation and metabolic homeostasis (*17*), showed substantially higher enrichment than the other transcription factors. Overall, the results indicate that altered gene expression, particularly in muscle-related pathways and energy metabolism, occurs before the onset of symptoms in ALS.

### Gene clustering and ontological assignment highlights coordinated transcriptional changes in key molecular pathways

We used weighted gene co-expression network analysis (WGCNA) to identify modules of co-expressed genes in the muscles of PRE-ALS individuals, assuming that genes exhibiting correlated expression changes across conditions likely participate in shared biological processes and thus belong to the same functional group (*18*). This analysis revealed eight gene clusters (Fig. 2A), where genes within each cluster exhibited similar expression trends (Fig. S1). We used an adjacency matrix and topological overlap measure to estimate gene connectivity within the network, demonstrating significant connections in the modules.

**Figure 2.**
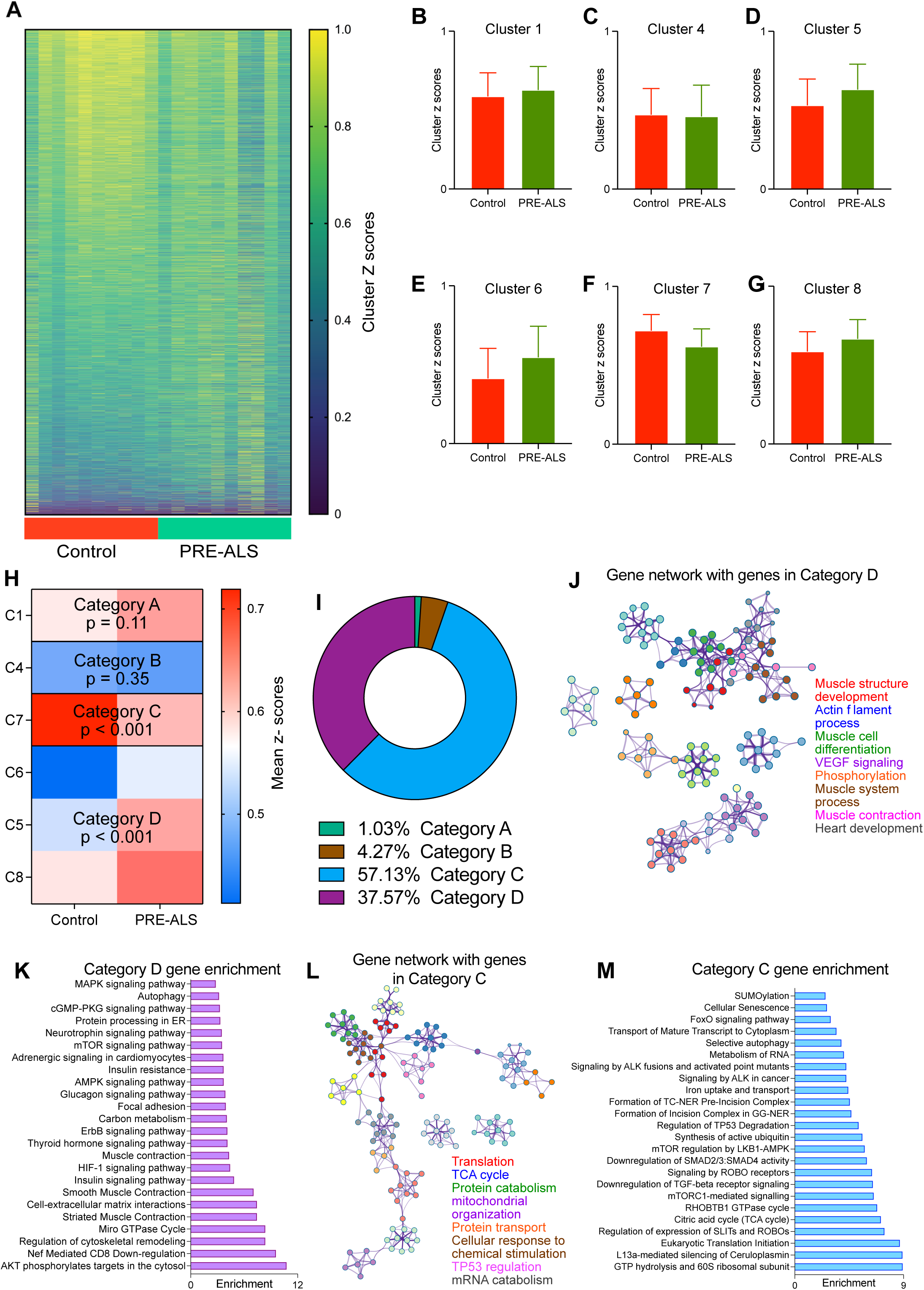
Clustering of genes indicates key pathways segregated into two broad categories. (A) Gene expression correlation heatmap revealing distinct clustering patterns in control and PRE-ALS participants. Clustering patterns were generated using weighted gene co-expression network analysis (WGCNA). (B-G) Mean z-scores of the clusters of genes identified as cluster 1(B) 4 (C), 5 (D), 6 (E), 7 (F) and 8 (G) among control and PRE-ALS participants. (H) Heatmap of the categories classified from clusters 1, 4, 5, 6, 7 and 8 based on their global trends in the mean z-scores among control and PRE-ALS participants. Category A consists of genes in cluster 1 (p = 0.11). Category B consists of genes in cluster 4 (p = 0.35). Category C consists of genes in cluster 7 (p < 0.001). Category D consists of genes in clusters 5, 6 and 8 (p < 0.001). (I) Pie chart demonstrating the percentage of genes representative of each category. (J) Gene network representation generated from genes in category D. (K) Pathway enrichment and key pathways impacted by the genes in category D. (L) Gene network representation generated from genes in category C. (M) Pathway enrichment and key pathways impacted by the genes in category C.

We assessed Z-score trends of the entire gene set without an initial p-value filter to capture the cumulative effect of subtle, coordinated changes in gene expression. To assess overall expression trends, we compared the average normalized Z-score of each cluster, excluding smaller clusters with fewer than 50 genes to ensure robust results (Fig. 2B–G) - for example, clusters 2 and 3 were excluded due to their small size (14 and 31 genes, respectively). A Z-score trend reflects a cluster’s collective expression becoming more or less active relative to the gene set average, revealing a directional shift in expression. Based on the statistical significance and directionality of these trends, we grouped the clusters into categories (Fig. 2H). Categories C (cluster C7) and D (clusters C5, C6, and C8) particularly stood out, accounting for approximately 57% and 38% of the statistically significant genes, respectively (Fig. 2I). The remaining clusters, C1 and C4, showed no significant changes.

Gene networks in category D are involved in muscle structure development and differentiation, actin cytoskeleton, and VEGF signaling (Fig. 2J). Further enrichment analysis suggested involvement in MAPK, AKT phosphorylation, AMPK and insulin signaling pathways (Fig. 2K). Genes in category C were enriched for metabolic pathways, including the TCA cycle, protein transport and catabolism, mRNA catabolism, etc. (Fig. 2L–M).

These findings reveal coordinated shifts in muscle gene activity, pointing to early dysregulation of structural and metabolic pathways in the muscle of asymptomatic ALS gene carriers.

### Muscle metabolic dysfunction arises from the interaction of multiple dysregulated pathways

Initial analysis revealed key regulatory relationships and influential gene cluster groups (categories) within the transcriptional landscape. We identified 2,105 transcriptionally relevant genes by selecting the top 30% most variable genes with a p-value <0.05. To assess the functional importance of gene clusters, we employed a topological analysis framework, using a negative degree change to identify clusters with the highest network impact (Fig. 3A, left).

**Figure 3.**
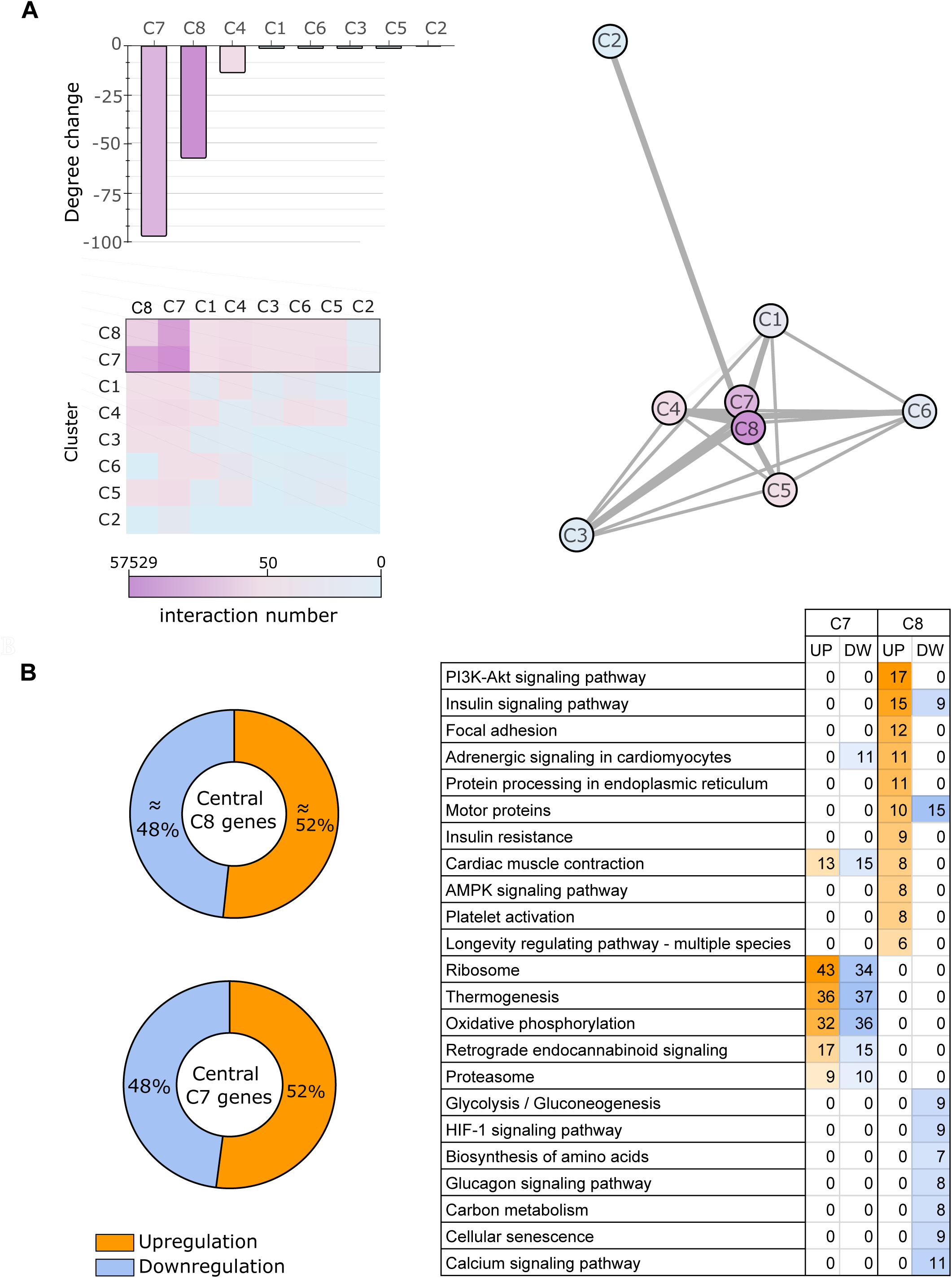
Identification of central clusters related to the observed changes within the dataset. (A) The network’s cluster distribution and interactions, with C7 and C8 identified as the most influential clusters based on their negative degree change indicating their greater impact on the network’s topology. (B) A total of 367 and 234 central genes were identified in C7 and C8, respectively. Central genes were identified based on their degree change, similarly to clusters. The accompanying table details the number of up- and downregulated genes (padj<0.05) within specific pathways for each cluster.

Clusters C7 and C8 emerged as the most influential, occupying central positions and driving the network’s configuration, with C7 exhibiting a more core, hub-like centrality (Fig. 3A, right). Both clusters contained a near-even mix of up- and downregulated genes compared to controls (∼48% down, ∼52% up; Fig. 3B, left). Cluster C8 showed a stark dichotomy between upregulated and downregulated pathways, suggesting distinct subnetworks. Upregulated genes were enriched in pathways like PI3K-Akt, insulin, and AMPK signaling, while downregulated genes were associated with glycolysis/gluconeogenesis, glucagon signaling, and carbon metabolism (Fig. 3B, right). The insulin signaling pathway was unique in its enrichment with both up- and downregulated genes, with the latter showing overlap with other downregulated pathways (Tables S3, S4). Key genes driving metabolic dysregulation were also identified. Downregulation of *CaMK2B*, a central node in contraction-induced glucose uptake, suggests impaired muscle glucose handling (*19*). Dysregulation of *HK1* and *PHKG1*, which encodes for critical enzymes in glucose metabolism and glycogenolysis respectively, further supports altered glucose handling (*20*). *FLOT1*, a lipid raft protein associated with insulin signaling, may contribute to metabolic dysfunction (*21*). *MLXIP* and *MLXIPL* genes, involved in the regulation of genes in response to cellular glucose levels (*22*), were also dysregulated. Finally, *SREBF1* (encoding transcription factors sterol regulatory element binding proteins SREBP1a and 1c), and *ACACB* (encoding acetyl-CoA carboxylase 2, ACC2), both involved in fatty acid metabolism (*23, 24*), were identified as dysregulated. Collectively, these findings point to a complex interplay of dysregulated pathways underlying muscle metabolic dysfunction.

### Insulin and AMPK signaling are drivers of the dysregulated muscle transcriptome

Network-based analysis was employed to identify key genes and pathways from enriched gene clusters C7 and C8 within the presymptomatic stage. We used a combined metric of three centrality measures - degree, betweenness, and eigenvector - to capture different aspects of nodal influence. Degree centrality measures direct connections, betweenness centrality identifies nodes on critical network paths, and eigenvector centrality weights a node’s importance by its neighbors’ connectivity (*25*). This approach allowed us to classify nodes based on their network role: highly active drivers (high values for all three measures), pathway regulators (high degree and eigenvector), and critical connectors (high betweenness and eigenvector) (Fig. 4A and 4B). Genes were ranked by their centrality, with negative degree changes indicating increased influence. Enrichment analysis of centrally-ranked genes in cluster C8 identified key pathways (Fig. 4C). A subsequent pathway interaction graph revealed insulin signaling as a highly influential pathway, consistently demonstrating high centrality scores in C8 (Fig. 4C and 4D). The AMPK signaling and thermogenesis pathways were identified as essential mediators, while MAPK and thyroid hormone signaling appeared to be regulatory factors (Fig. 4D).

**Figure 4.**
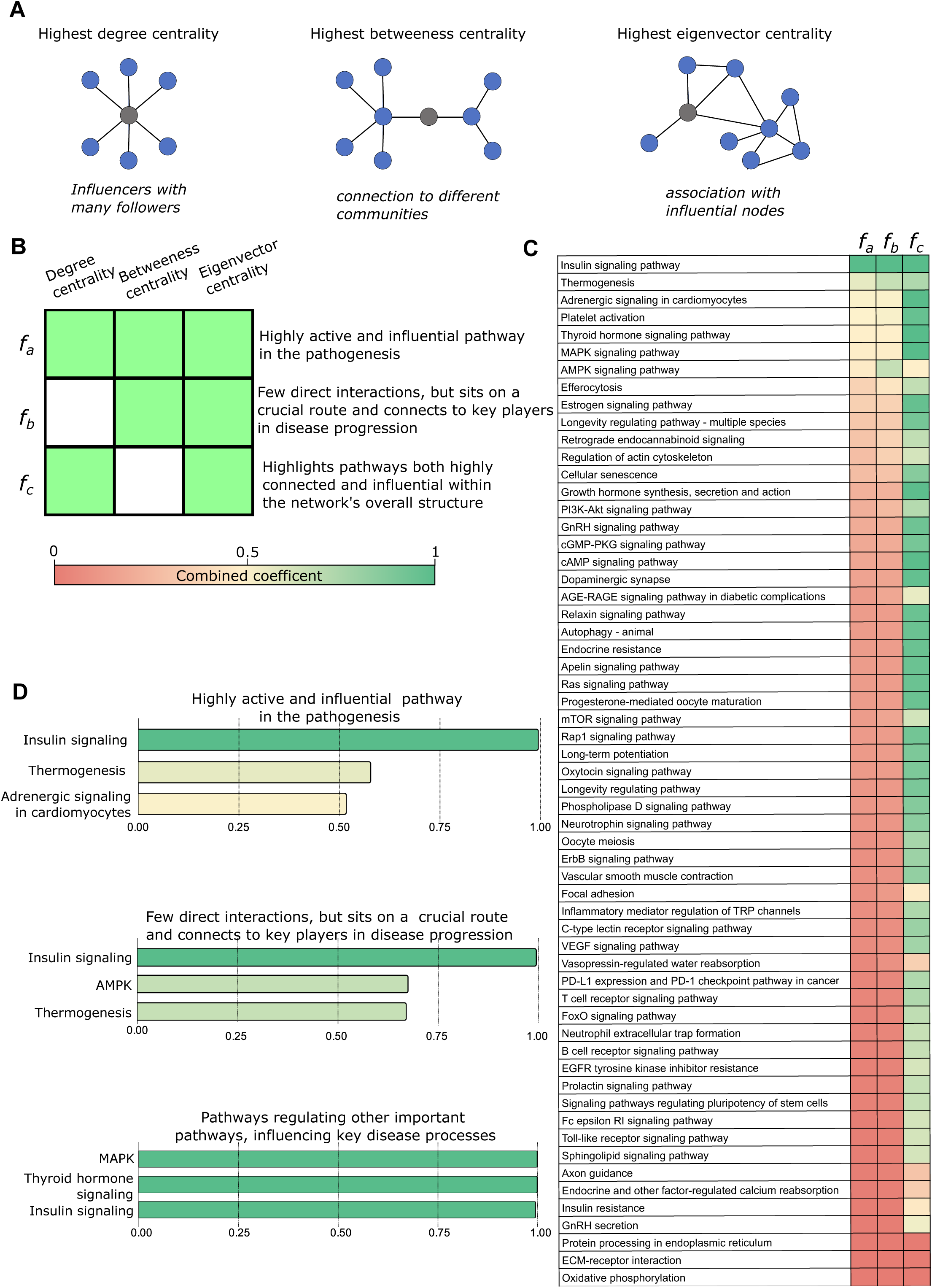
Unraveling functional roles of central gene clusters: Network inspired analysis of enriched pathways of cluster 8. (A) Three key network centrality measures quantify a node’s relative importance: degree centrality (number of connections), betweenness centrality (bridging two distinct subclusters), and eigenvector centrality (connectivity to important nodes). Each metric captures a distinct aspect of a node’s connectivity and influence. (B) Normalized measures of eigenvector, degree, and betweenness centrality (range [0, 1]) were combined to create three coefficients (fa, fb, fc). fa (all three centralities) identifies highly active and influential pathways. fb (betweenness-eigenvector) highlights critical connector pathways that facilitate signal transmission. fc (degree-eigenvector) highlights pathways that are both highly connected and influential. Higher coefficient values indicate increased prominence of the respective node characteristics. (C) Enriched pathways from an ordered list of central genes in C8 and their associated coefficients. Genes were ranked by their overall impact on the cluster network: a more negative degree change indicates a more central gene. Pathway interaction was identified and constructed through enrichment analysis of this ranked gene list. (D) The top three C8 pathways scored by each coefficient. The design of the first coefficient, results in its high values coinciding with high values for the second and third coefficients.

A similar analysis of cluster C7 revealed that the thermogenesis pathway exhibited high centralities, analogous to the insulin signaling pathway in C8 (Fig. 5A). Although the “metabolic pathways” score was modest, its high rank in the combined centrality analysis suggested a potential role in energy metabolism. Further analysis of this pathway in C7 revealed the TCA cycle and related processes (respiratory electron transport and ATP synthesis) as highly enriched (Fig S2 and associated data). Following transcriptional motif enrichment analysis, we subsequently identified estrogen-related receptor alpha, a key transcription factor responsible for the transcriptional control of metabolic genes, which is reported to be significantly influenced by both AMPK and insulin signaling pathways, with downstream effects impacting the TCA cycle, oxidative phosphorylation, and post-translational modifications (*26*), as predominantly enriched (Fig. S2). Our results also suggest that autophagy and c-type lectin receptor signaling pathways might act as regulatory factors within the information transmission network (Fig. 5B).

**Figure 5.**
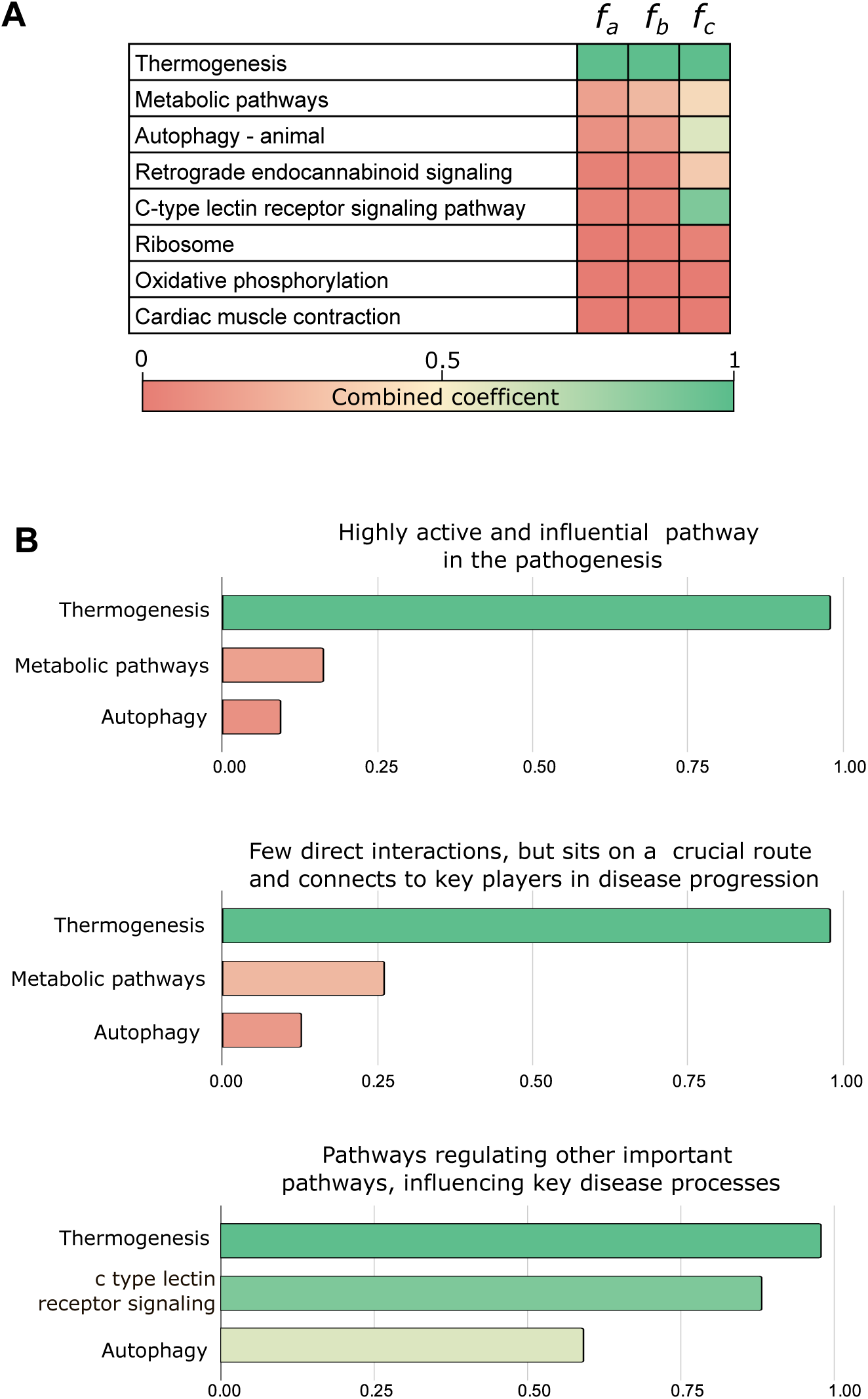
Network inspired analysis of enriched pathways of cluster 7. (A) Enriched pathways of the ordered list of central genes from cluster 7 and their associated coefficients. (B) Top three scored pathways based on the coefficient values. Thermogenesis (i.e. contraction-mediated heat production) and metabolic pathways (mainly represented by the TCA cycle -Fig S2), indicate that cluster 7 seems to be highly involved in energy expenditure and metabolism.

Our results suggest that gene cluster C8 may exert both compensatory and detrimental effects, potentially transmitted to the more central C7. We identified the thermogenesis pathway as a key conduit for information flow between clusters C7 and C8, suggesting that C8 may be the primary site for the manifestation of significant functional changes. In addition to thermogenesis, which links the two clusters, the AMPK pathway within the C8 cluster also represents a critical pathway for mediating transmission.

Overall, our analysis indicates that insulin signaling, the AMPK signaling pathway, and metabolic regulation are the primary contributors to the observed dysregulated transcriptome, with insulin signaling emerging as a central driver and AMPK signaling as a key regulator. The TCA cycle and thermogenesis pathways likely represent downstream responses rather than primary drivers of these transcriptional alterations.

### Systems analysis identifies presymptomatic gene expression pattern changes, unmasking a potential compensatory molecular mechanism

Our initial network analysis of differentially expressed genes (DEGs) focused on the top 30% most variable genes (p<0.05) between the PRE-ALS and control groups. Using enriched pathway and topological network analysis, we identified key biological processes dysregulated in PRE-ALS participants. However, the PRE-ALS cohort is heterogeneous, most probably including individuals at various stages of pathogenesis before phenoconversion. Relying solely on a strict DESeq2 p-value threshold may obscure subtle but biologically important gene expression changes present in a subset of individuals who will eventually develop ALS. Therefore, we expanded our analysis beyond the initial threshold to capture these signals. We highlighted additional DEGs, with their nominal and adjusted p-values that were relevant to our key pathways, providing a more comprehensive view of the biological landscape.

Our transcriptomic analyses revealed the insulin signaling pathway, AMPK signaling, and the TCA cycle as central to the associated transcriptomics alterations in the PRE-ALS participants. Since the mechanistic target of rapamycin (mTOR) signaling acts as a critical convergence point integrating inputs from both AMPK and insulin inputs (*27*), we investigated its role in the observed changes. We assessed correlations between specific model components by investigating their transcriptomic profiles. Our approach quantified the degree of coordinated pathway activation by determining the proportion of downstream gene expression variance attributable to upstream gene expression using correlation coefficients (r^2). For branches regulated primarily at the post-translational level (blue branches, Fig 6A), we inferred activity from downstream transcriptional changes.

**Figure 6.**
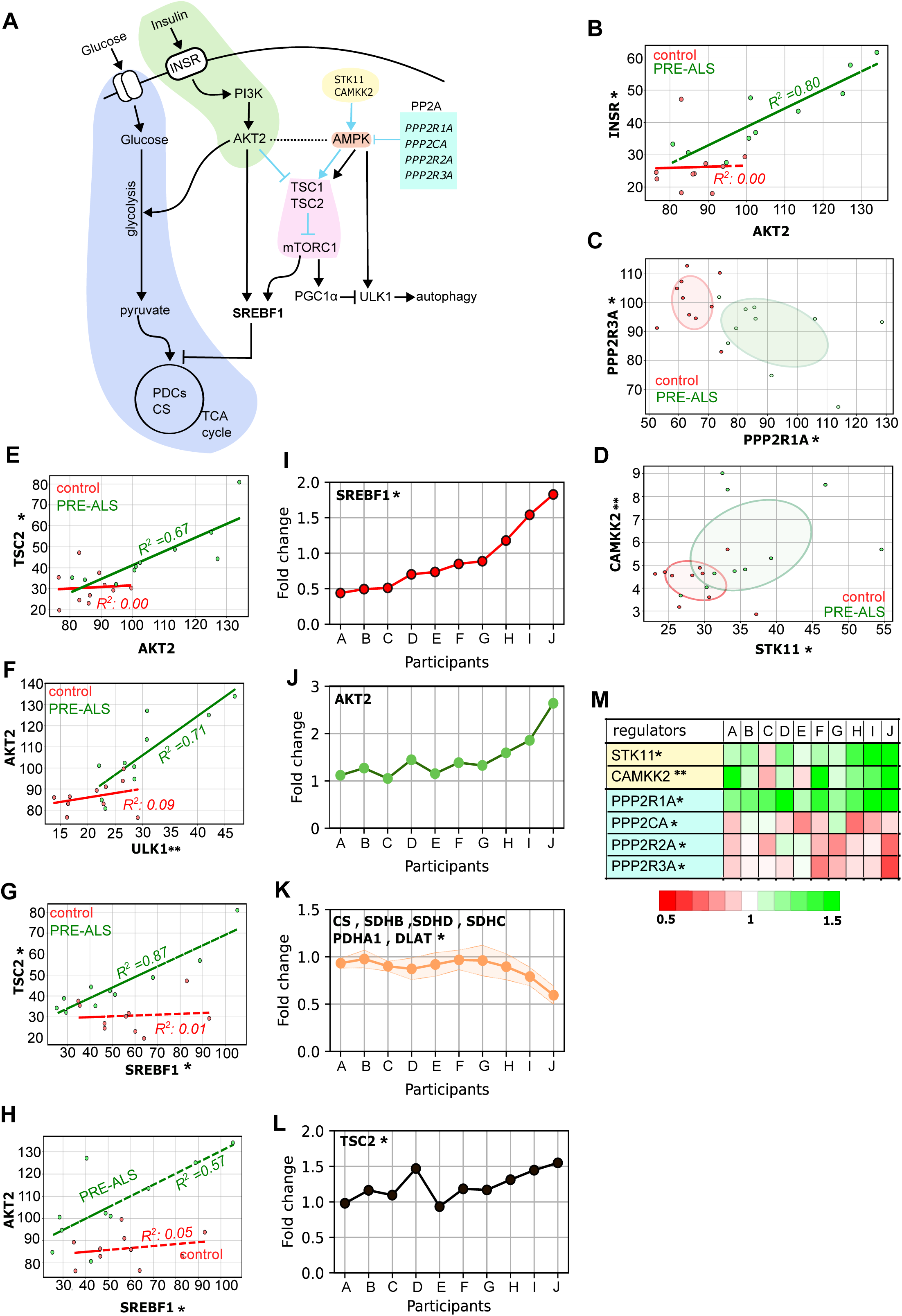
Pre-symptomatic dysregulated muscle pathways and hypothetical dynamical landscape illustrating the proposed progression of ALS along an imaginary time axis. (A) Interplay among the insulin signaling pathway, AMPK signaling, and the mTOR pathway, alongside other key regulatory components identified as central to transcriptomic alterations in PRE-ALS participants. Branches primarily regulated at the post-translational level are in blue. (B-M) Based on the DESeq2 group-level analysis, (*) indicates a nominal p-value < 0.05 but a non-significant adjusted p-value, while (**) denotes a result that is non-significant by both measures. For branches primarily regulated at the post-translational level activity levels were indirectly inferred through analyses of their downstream transcriptional targets. (B) Upstream and downstream insulin signaling components show increased activation relative to controls, with component correlations consistent with an active pathway. (C-D) Changes in PP2A subunits (*PPP2R1A* and *PPP2R3A*) and activation of upstream regulators (*STK11* and *CAMKK2*) suggest a shift in AMPK signaling. (E-H) Gene correlation and R² analysis exposes network dysregulation of the TSC2-mTOR regulatory axis, leading to elevated *SREBF1* and autophagy pathway signaling, particularly through ULK1. (I-M) Gene expression profiles in the PRE-ALS group. Participants are ordered from lowest to highest level of *SREBF1* expression. Fold change values are normalized against the control group. Collective expression of TCA cycle-associated genes is displayed, with the dashed region representing the standard deviation of the mean. Individual plots are shown for other key genes in the model.

We observed significant differential expression of key insulin signaling components, INSR and AKT2 (padj<0.05), with consistent changes across participant groups relative to the control (Fig. 6B). These findings are particularly relevant given that, when activated, INSR can initiate the PI3K-AKT cascade (*28*). Namely, within the PRE-ALS group, a strong positive linear correlation (r²≈0.8) between INSR and AKT2 expression was observed, suggesting their tight coupling during the presymptomatic stage. This relationship was absent in the control group and indicates potential progression towards altered metabolic regulation.

To further understand the regulatory landscape of these pathways, we analyzed the expression of AMPK regulators -STK11, CAMKK2 (known activators) (*29, 30*) and key regulators of the protein phosphatase PP2A (known repressor) (*29, 31*).

Our analysis focused on statistically significant subunits of PP2A given their extensive isoform diversity (*32*). We identified significant differential expressions (padj<0.05) of *PPP2R1A* (scaffold subunit) and *PPP2R3A* (regulatory subunit) between the PRE-ALS and control groups (Fig. 6C). The observed upregulation of *PPP2R1A* and downregulation of *PPP2R3A* suggest a potential increase in PP2A complex formation with compromised regulatory control. Furthermore, while not statistically significant, we also observed a trend of upregulation for the AMPK activators, *STK11* and *CAMKK2*, in some of the PRE-ALS participants (Fig. 6D).

We observed a positive mRNA level correlation between *TSC2* and *AKT2* that was both significant and unconventional (Fig 6E). The observed increase in *TSC2* mRNA levels in specific PRE-ALS participants may be driven by AMPK activation, as AMPK is known to transcriptionally upregulate *TSC2* expression (*33*). Consistent with the expected inhibition of mTORC1 by elevated TSC2 levels (*34*), our data shows that increased *TSC2* transcripts correlated with reduced PGC-1α mRNA (*35*) (Supplementary Information Datafile04). This decline in *PGC-1α* is known to be linked to the induction of autophagy (*36*). In agreement, *ULK1* levels also correlated with the rise in *AKT2* (and consequently TSC2) in specific participants (Figure 6F), suggesting early activation of autophagy, a process known to be promoted by AMPK and inhibited by mTOR (*37*), an observation once again consistent with transcriptional changes observed in our dataset.

Contrary to the expected inhibitory effect of TSC2 on mTORC1 signaling and downstream *SREBF1* expression (*34, 35*), *TSC2* mRNA levels positively correlated with *SREBF1* mRNA in our dataset (Fig. 6G). This finding, supported by a strong *AKT2-SREBF1* expression correlation (Fig. 6H), suggests activation of an mTOR-independent *SREBF1* pathway driven by AKT2 (*38*).

### Gene expression stratification validates the presymptomatic molecular signatures identified and predicts phenoconversion risk in statistically robust analysis

We next focused on analyzing individual fold changes relative to the control group, with PRE-ALS participants ordered based on the expression level of *SREBF1* (Fig. 6I). This approach allowed us to approximate the ‘temporal evolution’ of gene expression, revealing the dynamic processes that may precede symptom onset, and overcoming the limitations of group-level comparisons.

Using this participant order, we tracked the expression of other model components, including the mTOR axis (*AKT2, TSC2*), AMPK regulators (PP2A subunits, *STK11, CAMKK2*), the pyruvate dehydrogenase complex (*PDHA1, DLAT*), and key TCA cycle components (*CS, SDHB, SDHD, and SDHC*) (Fig. 6J-M). Given that *SREBF1* functions as a key metabolic gatekeeper that coordinates metabolic reprogramming (*39*), the same participants with the highest *SREBF1* dysregulation also exhibited the highest dysregulation in these other pathways, predicting them to be closest to phenoconversion (Fig. 6I-L). To validate this observed temporal progression and confirm that it reflects increasing network disruption, we employed a monotonic trend test on genes from key pathways: insulin, AMPK signaling, and TCA metabolism (Fig. 7). Genes were categorized by their functional association with these processes, including pathway membership, specificity, and regulatory functions into:

**Figure 7.**
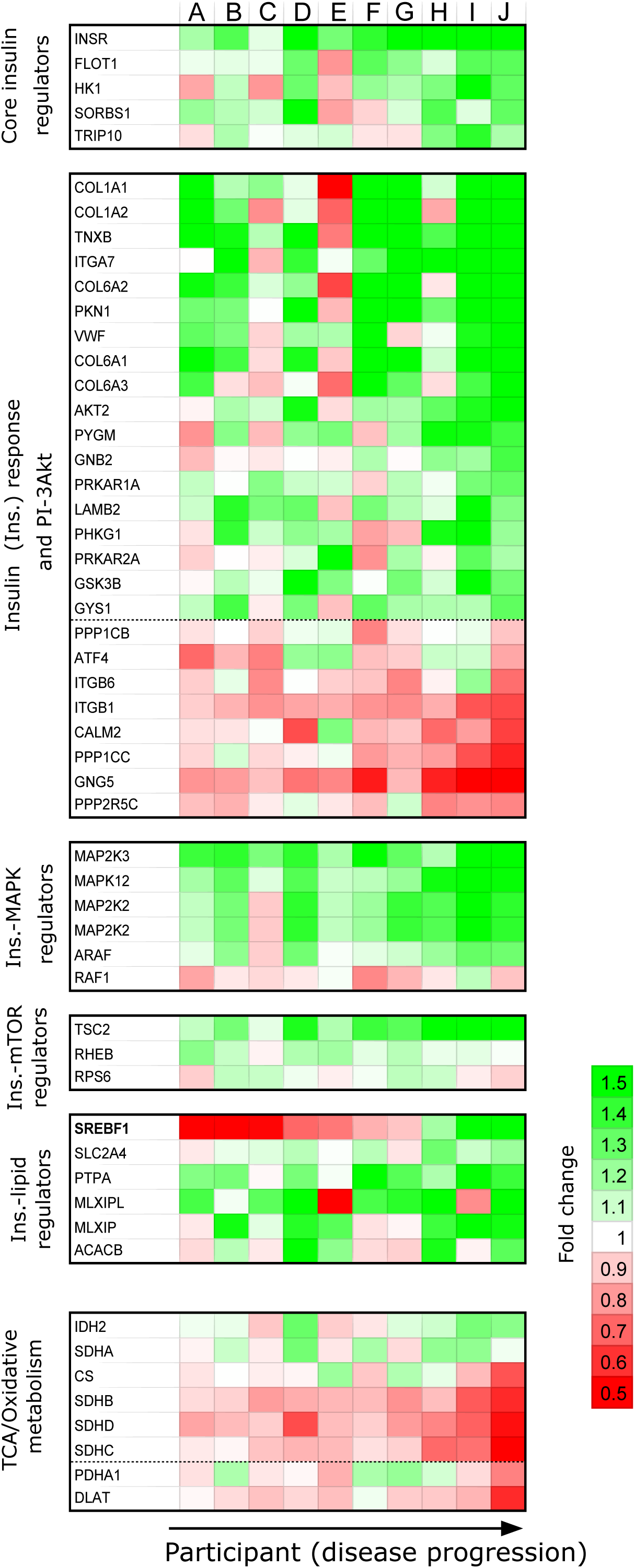
Gene expression map of key genes and pathways validates the prediction set by the proposed model of presymptomatic ALS and demonstrates statistical optimality. Participant stratification based on their likelihood of developing clinically-manifest ALS. Gene expression data, shown as a fold change relative to controls, illustrates the patterns associated with disease probability. This analysis uses the same key pathways and processes identified by the proposed model. Participants are arranged from the lowest (A) to the highest (J) probability of developing clinically-manifest disease, with genes grouped by their functional associations: Groups I - VI encompassing different signaling pathways - core insulin regulators (I), insulin response and Akt (IIa/b), insulin-MAPK regulators (III), Insulin-mTOR regulators (IV), Insulin - lipid regulators (V), and TCA/oxidative metabolism (VI). All genes originated from central clusters C8 and C7 with the exceptions PDHA1 and DLAT, which were included due to their close proximity and association with the other genes within the VI group.

I: Genes involved in the insulin signaling pathway, including mediators, downstream receptor effectors, and those regulating glucose metabolism.

IIa, IIb: Genes that translate AKT signals into metabolic actions, regulating targets PI3K-AKT pathway mediators and other downstream effectors of AKT.

III: Genes linking the MAPK and insulin signaling pathways, which influence glucose uptake, muscle growth, and cellular stress response.

IV: Insulin responsive mTORC1 regulators (TSC2, RHEB2) and effector RPS6, activated by AKT signaling.

V: Insulin-responsive metabolic regulators involved in lipogenesis and cholesterol synthesis.

VI: Cellular metabolism genes, primarily encoding TCA cycle enzymes. All genes originated from central clusters C8 and C7, with the exceptions of PDHA1 and DLAT, which were included due to their close association.

To ensure statistical robustness of participant ordering, we evaluated a permutation of 100,000 different ways of ordering participants based on their gene expression profiles (Fig. S3). Only three permutations met this stringent criterion (Table S5), and all three accurately matched our hypothesized order. Specifically, participant “A” consistently ranked as lowest risk, and “J” was invariably placed at the highest risk, demonstrating the ability to distinguish between asymptomatic individuals least and most susceptible to developing clinically-manifest ALS. Analysis of expression patterns and participant ordering was supported by a greater than 80% increase in serum NfL levels in the participant classified as highest-risk during follow-up.

To examine pathway behavior, we quantified progressive monotonic pathway perturbation across ordered participants using an ’accumulator’ metric, which tracks gene expression trends by adding or subtracting 0.1 for higher or lower expression, respectively (Fig. S4). Our findings suggest that dysregulated pathways in PRE-ALS individuals are highly interconnected with a significant overlap in genetic components. Across accumulator thresholds, oxidative phosphorylation remained consistently downregulated, whereas TCA, AMPK, and insulin signaling pathways showed clear expression shifts only under the strictest criterion. Notably, participants predicted to be closer to clinical onset (I and J) showed higher standard deviations in gene expression, suggesting increased variability and pathway dysregulation with a potential breakdown of normal regulatory mechanisms.

## DISCUSSION

In this study, we investigated muscle gene expression patterns in ten asymptomatic - defined as the absence of any clinical signs or symptoms of overt disease (*40*) - carriers of ALS gene mutations. We report for the first time that skeletal muscle from asymptomatic ALS mutation carriers undergoes a complex metabolic reprogramming, detectable far before the expected disease onset.

Our first major finding is that specific modifications of gene expression profiles occur in asymptomatic ALS mutation carriers, preceding detectable changes in the known phenoconversion biomarkers assessed in this study. We revealed that the genes within the two most influential gene clusters were linked to the insulin signaling pathway, thermogenesis pathway, AMPK signaling pathway, and TCA cycle. These pathways act as the main contributors to the dysregulated muscle transcriptome. Moreover, the insulin and AMPK signaling pathways are likely co-activated in the skeletal muscle of asymptomatic ALS mutation carriers. In resting conditions, insulin signaling is the main anabolic pathway that triggers the storage of energy-rich molecules in muscle. The insulin pathway promotes glucose uptake and glycogen storage in skeletal muscle (*41*). It also facilitates fatty acid uptake and their storage as triglycerides, while simultaneously stimulating protein synthesis and suppressing protein degradation through activation of mTORC1 (*41*). Thus, beyond its well-known role in increasing glucose uptake to feed glycolysis, insulin strongly stimulates glycogen, lipid, and protein synthesis. By contrast, AMPK signaling is activated in the skeletal muscle in situations of energy stress (i.e., energy intake deficits or increased energy demand) or nutrient deprivation (*42*), leading to an increase in cellular AMP/ATP and ADP/ATP ratios (*43*). It acts as a metabolic switch, turning off anabolic processes and favoring glucose and fatty acid catabolism (*44*). Therefore, the simultaneous activation of potent anabolic and catabolic pathways in the muscle of asymptomatic ALS gene carriers is unexpected and is likely to severely disrupt muscle physiology. To our knowledge, the post-exercise recovery is the only circumstance in which this type of combined activation could be recorded. Indeed, depending on exercise intensity and duration, different AMPK complexes are activated in skeletal muscle (*45*). Short-duration high-intensity and prolonged moderate-intensity exercise have both been shown to increase AMPK activity and ACC2 phosphorylation, leading to progressive increase in fatty acid oxidation (*46, 47*). After exercise/muscle contraction, AMPK signaling remains activated, in parallel to insulin signaling activation, which feeds muscle cells through the increase of nutrient intake. This co-activation enhances glucose uptake through additive effects on GLUT4 translocation via converging phosphorylation of TBC1D4 by both AKT and AMPK (*48*). Overall, the co-activation of insulin and AMPK pathways in the muscle of asymptomatic ALS gene carriers could lead to increased muscle insulin sensitivity together with a progressive increased reliance on fatty acid oxidation (*49*).

When analyzing heterogeneous populations, such as those in the PRE-ALS cohort, the DESeq2 pipeline presents limitations. Its negative binomial model, while effective for identifying average gene expression trends, may not fully capture substantial inter-individual variability. This can lead to misinterpretation where a statistically significant downregulation represents a systematic shift in the average expression rather than a consistent decrease across all individuals. For instance, a small subset of individuals showing an upward trend in gene expression might be masked by a more widespread downregulation across the group.

Using a targeted approach focusing on the fold change of each participant relative to the control group in the previously established critical pathways, our data suggest that metabolic reprogramming in muscle unfolds progressively during the transition to phenoconversion. Analysis of pathway dysregulation during the transition between control and PRE-ALS participants revealed a progressive increase in expression of *SREBF1*, a master transcriptional regulator of lipid synthesis, in parallel with progressive activation of AMPK and insulin signaling pathways. Notably, this observation aligns with our finding that *SREBF1* exhibited an mTOR-independent activation pathway driven by AKT2. *SREBPF1* encodes SREBP1a and 1c transcription factors – SREBP1c being the predominant isoform in skeletal muscle (*50*) – which are key actors of the regulation of genes related to lipid metabolism (*24*). SREBP1c connects insulin signaling to lipid metabolism (*51*) and, in the presence of glucose, activates transcription of muscle genes required for fatty acid synthesis and oxidation, including *ACACB* (*52*).

Taken together, our results thus point toward a progressive activation of fatty acid synthesis and oxidation in skeletal muscle, mediated by increased activation of insulin signaling, during the transition from the presymptomatic phase to motor onset. Moreover, the chronic activation of AMPK in skeletal muscle is expected to enhance muscle fiber oxidative capacity by promoting mitochondrial biogenesis (*53, 54*), and drives a transition from fast (glycolytic) to slow (oxidative) myofiber programs (*55*). In resting muscle, AMPK activation further increases fatty acid oxidation through allosteric regulation of CPT-1 and transcriptional control of *PPARα* and *PGC-1* (*56*). These findings strongly suggest that concomitant dysregulation of insulin and AMPK signaling in the muscle of asymptomatic ALS gene carriers triggers an early shift toward oxidative metabolism - a hallmark of symptomatic ALS muscle - indicating that this reprogramming is likely an important contributor to disease pathogenesis. Strikingly, in PRE-ALS individuals identified as being closest to phenoconversion in our model (i.e. participants I and J), we observed a collapse in expression of the main components of PDH complex - the key gateway linking glycolysis to the TCA cycle (*57*) - as well as of TCA cycle components. This is likely to result in major metabolic stress through a drop in ATP production via oxidative phosphorylation. Overall, our model suggests that, in the muscle of ALS mutation carriers, the presymptomatic stage is characterized by progressive increase in muscle fiber oxidative capacity together with a progressive increase of fatty acid synthesis, oxidation and reliance. As phenoconversion approaches, oxidative phosphorylation efficiency in skeletal muscle appears reduced, potentially indicating diminished ATP-generating capacity.

Importantly, our data indicate that profound muscle metabolic reprogramming can be detected in asymptomatic ALS gene carriers before changes in established biomarkers - such as MUNIX or blood NfL levels - which are suggestive of the proximity of motor onset, become apparent. MUNIX provides an indirect measure of motor neuron loss, making it especially valuable for monitoring disease progression in ALS (*58*). In asymptomatic SOD1 mutation carriers, an abrupt reduction in motor unit number can be detected only in the late presymptomatic phase (*59*), with no corresponding data for C9ORF72 carriers. After motor onset, in ALS patients, a marked decline in MUNIX can be detected up to 12 months before overt clinical muscle weakness (*60*). In our study, assessment of MUNIX score showed no difference between PRE-ALS participants and controls at baseline, and no significant change was detected between baseline and the 18-month follow-up visit. Blood NfL is a biomarker of the aggressiveness of the neurodegenerative process, which has been proposed as a biomarker for predicting the approach (within a few months to up to 5 years) of phenoconversion from presymptomatic to clinically manifest disease in ALS mutation carriers (*3, 4*). Participant J, predicted by our model to be the closest to motor onset, had elevated blood NfL at baseline (i.e. above the range observed in controls of the same age, (*15*)), and NfL levels increased by over 80% after 18 months compared to baseline. No other participant in our cohort showed elevated NfL levels at both baseline and 18 months, accompanied by a >25% increase during follow-up, a threshold demonstrated in healthy subjects to exceed natural biological variability between consecutive measurements (*61*). This strongly suggested that, at the end of study follow-up, participant J was the most at risk of transitioning to clinically manifest ALS.

This study has limitations. Recent data in asymptomatic ALS mutation carriers suggest that presymptomatic changes in body composition and resting energy expenditure may be mutation-specific (*13*). While our results provide an initial mechanistic explanation for presymptomatic systemic metabolic alterations, the relatively small sample size of our study precluded subgroup analyses to determine whether some of the observed muscle metabolic dysregulations might be mutation-specific. In addition, as muscle biopsy is an invasive procedure, it was only performed at inclusion in each participant, and was not repeated during study follow-up. Thus, we could not assess longitudinal changes in muscle gene expression profiles at an individual level. A longer-term clinical follow-up of PRE-ALS participants would also have been valuable to monitor a potential transition from the pre-symptomatic to the symptomatic disease phase. Finally, the study was designed before the recent recognition of prodromal states within the presymptomatic phase, which may manifest as mild motor, cognitive, or behavioural impairment (*62*). At the end-of-study visit, needle EMG was performed only in two regions (two muscles in the upper limb and one in the lower limb) on the side used for MUNIX recording, thus not meeting the recently proposed research criteria for mild motor impairment (*63*). All participants underwent Mini Mental State Examination (MMSE) at inclusion, but longitudinal neuropsychological assessment was not conducted. Overall, our data do not allow us to determine whether the observed muscle metabolic phenotype would be specific to phenoconversion toward ALS or might also occur in FTD.

## Conclusion

This study provides unexpected data by identifying major metabolic reprogramming in the skeletal muscle of asymptomatic ALS gene carriers, starting long before the onset of motor symptoms. These findings emphasize the critical role of energy homeostasis dysregulation as an early pathophysiological event in ALS. Further studies are now necessary to delineate the timing of systemic consequences resulting from these muscle metabolism abnormalities, ultimately paving the way for the development of novel early biomarkers of phenoconversion. Evidence that muscle metabolic reprogramming occurs in ALS from the presymptomatic stage highlights its potential role in disease progression and supports the rationale for future therapies targeting muscle metabolism.

## MATERIALS AND METHODS

### Participants

Ten asymptomatic individuals with a mutation in one of the two major ALS-associated genes - C9ORF72 and SOD1 – were included in the PRE-ALS study (Presymptomatic Neuromuscular Junction Defects and Compensatory Mechanisms in ALS, NCT03573466) between May 2019 and December 2021. All participants were recruited from the Pitié-Salpêtrière ALS Expert Centre and Reference Centre for Rare Dementia. All participants had been previously diagnosed on the basis of presymptomatic DNA tests performed in the context of familial ALS linked to SOD1 or C9ORF72 expansion or familial frontotemporal dementia linked to C9ORF72 expansion. At the time of enrollment, participants qualified as “asymptomatic” with no symptoms suggestive of ALS or cognitive impairment, and no signs of motor neuron disease at neurological examination performed by an expert neurologist of the team (MDMA, FS or GB). All participants underwent an open biopsy of the deltoid muscle at enrollment.

PRE-ALS participants were subsequently followed up every 6 months for 18 months, including a clinical interview, a neurological assessment and electrophysiological recordings. At the end of the study, participants resumed their routine medical monitoring with instructions to contact our team if any new symptoms occurred. The PRE-ALS study was sponsored by Assistance Publique - Hôpitaux de Paris and approved by the ethics committee (IDRCB number 2018-A00573-52).

For normal control samples, 10 deltoid biopsy specimens obtained from subjects considered free of any neuromuscular disorder after morphological and histochemical examinations were collected from the Biological Resource Platform of the Henri Mondor University Hospital.

All participants provided written informed consent consistent with institutional guidelines.

### Electrophysiological recording for Motor Unit Number Index determination

The number and size of motor units was evaluated using MUNIX at inclusion and every 6 months until the end of study follow-up, as described in the Supplementary Information. MUNIX and CMAP values recorded at baseline and 18 months were compared with those of 10 age- and sex-matched normal subjects previously recorded in the Neurophysiology Laboratory of the Pitié-Salpêtrière Hospital.

### Neurofilament light chain measurements

Plasma NfL was measured by Lumipulse at the baseline visit and at 18 months, and compared with values previously reported in healthy individuals of each age group (*15*) as described in the Supplementary Information.

### RNA sequencing

Total RNA from deltoid muscle was extracted as previously described (*14*). RNA library preparation was performed following the manufacturer’s recommendations (ILLUMINA Stranded total RNA prep ligation with Ribozero Plus) and as described in the Supplementary Information.

### Statistics

Statistical analysis was conducted with XLSTATS 2023 software (Addinsoft, Paris, France) and GraphPad Prism v10 software. The Mann-Whitney test or Wilcoxon signed rank test was applied when appropriate to compare continuous data. Fisher’s exact test was used to analyze categorical data. All statistics were two-tailed and the level of significance was set at p = 0.05. For the rest of the analytical pipelines, scripts and functions were generated using standard libraries in the Python programming language.

### Differential expression analysis using DESeq2

We used the DESeq2 package (*64*) to format a count matrix, where rows represent genes and columns are samples (control and PRE-ALS participants). Each entry in the matrix is the number of reads mapped to a specific gene. Before analysis, entries with zero reads in more than seven of ten samples per group were excluded. Genes with a false discovery rate (FDR) of less than 10% are considered significant. Similar methods have been employed elsewhere (*65–67*) and are explained in detail in the Supplementary Information.

### Weighted gene co-expression network analysis

WGCNA was used to study how groups of genes work together (*68*). The process involved selecting the top 30% of the most variable genes and constructing a network using a weighted Pearson’s correlation to create an adjacency matrix. Topological overlap measure was then applied to find genes with shared neighbors. By applying a soft-power threshold, the network was made to be “scale-free,” and hierarchical clustering was used to group genes into eight distinct modules based on their co-expression patterns. The expression trends within these modules were summarized using Z-scores, and Student’s t-test was performed to identify significant differences between the control and pre-ALS cohorts. WGCNA has been applied in clustering functional and co-expression-based gene or protein components in previous studies (*67, 69*) and is explained in detail in the Supplementary Information.

### Gene ontology analysis

Gene ontology and pathway enrichment analysis was performed on four gene pools using Metascape (*16*). We focused on gene ontology biological processes, cellular components, molecular functions, as well as KEGG and REACTOME pathways. The analysis parameters were set to a minimum overlap of 3, a p-value cutoff of 0.01, and a minimum enrichment of 1.5. Networks were mapped using STRING-db (https://string-db.org), with network edges defined by a high-confidence threshold (90%).

### Building interaction maps to study key pathways and regulators

To study key pathways and regulators, we constructed a bipartite network using the KEGG database (https://www.kegg.jp/kegg/ ) to map genes to their corresponding pathways. Nodes represented both genes and pathways, with edges connecting genes to their participating pathways. To focus on the most biologically relevant processes, we filtered the network to include only the top 30% of the most highly variable genes and those that were statistically significant and differentially expressed between the two groups (p<0.05). This step facilitated the identification of key biological processes and regulatory elements. Building upon this enriched gene-pathway network, we derived three distinct subnetworks to analyze specific interactions: a gene-gene network connecting genes with shared pathway membership, a pathway-pathway network connecting pathways that share genes, and a cluster-cluster network based on WGCNA to explore the interrelationships between gene co-expression modules.

### Assessment of key nodes within the network - identifying key clusters and gene ranking for functional relevance

To assess the importance of individual nodes, we employed a degree centrality-based metric. We evaluated each node’s influence on the overall network connectivity by systematically removing it and recalculating the network’s degree centrality. Nodes causing a greater decrease in average degree were considered more central and crucial for network integrity. This method was applied to both the cluster-cluster interaction map to identify key clusters and the gene-gene network to rank individual genes, defining those with a negative degree change as central genes.

### Network analysis and enriched pathway functional characterization of central genes

From this ranked list, only central genes were selected for an ordered enrichment analysis using g:Profiler (*70*). The resulting enriched pathways and their associated central genes were used to construct a new interaction graph. To characterize the role of each pathway, we applied three network centrality measures: degree (number of connections), betweenness (bridging nodes), and eigenvector (connectivity to important nodes). Each measure was normalized to a [0,1] range. Pathways were classified as disease drivers, key regulators, or critical connectors based on a composite score derived from these normalized measures.

## Supporting information

Supplemental informations

## Acknowledgements

The authors thank all the participants in the PRE-ALS study. The authors would also like to thank the team at the Neurosciences Clinical Investigation Centre at the Paris Brain Institute (Pitié-Salpêtrière Hospital, Paris), where the data used in this paper were collected, in particular Nadia Osman, Sabah Ait-Khelifa and Vanessa Brochard. The authors would also like to thank Mai Thao Bui and the team at the Risler Morphology Unit of the Pitié-Salpêtrière Hospital for invaluable technical assistance with muscle biopsies. Part of this work was carried out by the Paris Brain Institute Data Analysis Core (DAC) platform. (https://dac.institutducerveau.org/).

We gratefully acknowledge Emeline Cherchame for assistance with RNAseq data analysis. RNA quantification was performed at the CytoB2M core facility of BioMedTech Facilites (https://biomedicale.u-paris.fr/biomedtech-facilities/) INSERM US36 CNRS UAR2009 Université Paris Cité.

## Author Contributions

Conceptualization: GB

Methodology: LW, GB, FC

Investigation: CB, DM, DS, TL, FS, GB

Exploratory data analysis: DM, SS

Data modeling: DM, SS

Visualization: SS, DM

Supervision: SS, GB, LW

Writing - original draft: DM, SS, GB, LW

Writing - review & editing: all

## Data and materials availability

The RNA-seq datasets generated in this study will be deposited on Gene Expression Omnibus (NCBI) and made openly accessible upon publication. Other human-derived data will be made available from the corresponding author upon reasonable request, in compliance with applicable ethical and regulatory requirements.

## List of supplementary Materials

Materials and Methods

Supplementary Figures: Figs. S1 to S4

Supplementary Tables: Tables S1 to S5

Supplementary information for the 9 additional datafiles

Supplementary References

