## Supplemental informations for "Progressive muscle metabolic reprogramming in asymptomatic ALS gene mutation carriers"

Céline Buon et al.

Corresponding author:
Gaëlle Bruneteau,
Laure Weill,

**The pdf file includes**

Materials and Methods

Supplementary Figures: Figs. S1 to S4

Supplementary Tables: Table S1 to S5

Supplementary information for the 9 additional datafiles

Supplementary References

**Other Supplementary Material for this manuscript includes the following 9 datafiles**

Datafile01_diffexpr-results_counts.xlsx

Datafile02_fold_change.xlsx

Datafile03_Pathways and gene associations_supplementary figure 4.xlsxPathways and gene associations (Acc values).xlsx

Datafile04_PGC1alpha_deactivation - PGC1alpha.xlsx

Datafile05_preALS_supplementary_figure 3.xlsx

Datafile06_supplementary_figure 4,5.xlsx

Datafile07_metabolic_enrichment.xlsx

Datafile08_Ordered_Permutations.xlsx

Datafile09_Tested_Permutations.txt

**Material and methods**

**SM01. Electrophysiological recording for Motor Unit Number Index determination**

Surface electrophysiological recording was performed at inclusion to assess the number and size of motor units, and repeated every 6 months until the end of the PRE-ALS study follow-up at 18 months. The number of functional lower motor neurons was evaluated using Motor Unit Number Index (MUNIX), a fast, noninvasive, validated method to assess number and size of motor units in ALS patients (*1, 2*). MUNIX is calculated from the maximum compound muscle fiber action potential (CMAP) recorded in response to supramaximal electrical stimulation and voluntary surface electromyogram recordings associated with a series of submaximal voluntary muscle contractions. MUNIX was calculated for the abductor pollicis brevis, abductor digiti minimi, tibialis anterior, deltoid and trapezius muscles on one side. Measurements were obtained on the same side of the body throughout the study. In addition, at the end-of-study visit, concentric needle EMG was systematically performed to detect ongoing denervation changes (fibrillation potentials or positive sharp waves) in the first dorsal interosseous, TA and deltoid muscles, on the side used for MUNIX recording. A needle electrode evaluation could also be performed at any visit if clinical signs or symptoms raised suspicion of a prodromal state or phenoconversion.

The MUNIX sum-score and CMAP sum-score were calculated for each subject by adding the individual values of the 5 muscles tested. MUNIX and CMAP values recorded at baseline and 18 months were compared with those of 10 age- and sex-matched normal subjects previously recorded in the Neurophysiology Laboratory of the Pitié-Salpêtrière Hospital.

**SM02. Neurofilament light chain measurements**

As part of the study protocol, blood samples were collected into tubes containing EDTA at inclusion and after 18-month follow-up. Samples were centrifuged at 3500 rpm for 10 minutes at 4C aliquoted in polypropylene tubes and stored at -80C until use. The concentration of plasma NfL was measured using chemiluminescence enzyme immunoassay on Lumipulse G1200 (Fugirebio) according to the manufacturer's instructions. Plasma samples from participants were analyzed in a single run to ovoid the between run variability. Blood NfL concentrations determined in each PRE-ALS participant were compared with values previously reported in healthy individuals of each age group using SIMOA™ (*3*), a technique that has been shown to produce blood NfL results in high agreement with those obtained using Lumipulse™ (*4, 5*).

**SM03. RNA sequencing**

Total RNA from deltoid muscle was extracted using Tri Reagent (TR118, Molecular Research Center). 5 mm stainless steel beads (69989, Qiagen) were added to the Tri Reagent and extraction was performed in a tissue lyser. Then, total RNA was ethanol precipitated with 20 µg of glycogen (08-0111, Euromedex). RNA library preparations were made following the manufacturer's recommendations (ILLUMINA Stranded total RNA prep ligation with Ribozero Plus). Final samples of pooled library preparations were sequenced on ILLUMINA Novaseq 6000 with Ribozero Plus. Final samples of pooled library preparations were sequenced on ILLUMINA Novaseq 6000 with SP-200 cycles cartridge (2x800Millions of 100 bases reads), corresponding to 2x30Millions of reads per sample after demultiplexing.

The bioinformatics analyses were performed by the Data Analysis Core Facility (RRID:SCR_026138). The quality of raw sequencing data was evaluated with FastQC ((*6*) available online at: http://www.bioinformatics.babraham.ac.uk/projects/fastqc/). Poor-quality sequences were removed and adapters were trimmed with fastp using default parameters (Fastp: (*7*)). The Illumina DRAGEN bio-IT Platform (v3.10.4) was used for mapping the reads against the reference genome hg38, and for quantifying the reads based on GENCODE v.37. Library orientation, library composition, and coverage along transcripts were evaluated with Picard tools. Data visualizations were conducted with R shiny private software named Quby (https://dac.institutducerveau.org/). First, edgeR (*8*) was used to calculate normalized counts per million (CPM) (v3.38.0)

**SM04. Differential expression analysis using DESeq2**

*Preparing DESeq2 data class:* The DESeq2 package (*9*) in R BioConductor (http://www.bioconductor.org/packages/release/bioc/html/DESeq2.html) was used to format the count table into DESeq2 data class. The counts data was normalized using log transformation and normalized log transformation and used for differential expression analysis of the high-dimensional count data. The DESeq2 data class consists of a count matrix with rows corresponding to genes and columns denoting samples from pre-ALS participants cohorts. Each matrix entry indicates the number of reads unambiguously mapped to a gene in an experimental group.

*Preprocessing*: Data entries with reads equal to 0 for more than 7/10 samples per group-Control and pre-ALS participant cohort, were excluded from further analysis. For dimensional reduction and outlier identification, we performed a principal component analysis on the DESeq2 data class of count reads. Euclidean distance was used as a metric for determining orthogonal eigenvectors and eigenvalues. "*prcomp*" function in R was used, and variability explained by the eigenvectors was plotted as a biplot representation with the sequencing identity (control; pre-ALS participant cohort). The final biplot and heatmap of the covariance matrix shows the variability existing in this particular dataset.

*Differential expression using DESeq2:* The details of the DESeq2 pipeline are discussed in (*9*). Briefly, the DESeq2 package presents the data counts on the count matrix using a gamma-Poisson distribution with mean (normalized concentration of cDNA fragments from the gene in a sample). The size factors are determined by the median-of-ratios method. For each gene, a generalized linear model (GLM) is represented as a logarithmic fit, given by:

$$log_2 q_{ij} = \Sigma_r \; x_{jr} \beta_{ir}$$

where $x_{jr}$ are design matrix elements and $\beta_{jr}$ are the coefficients, with $j$ belonging to the pre-ALS participants cohort relative to the control for comparison between two groups. The GLM returns the overall expression strength of the gene, i.e., the log2 of the fold change (LFC) between the two groups compared (control versus pre-ALS participant cohort). Next, variability among replicates is modeled by a dispersion parameter $\alpha_i$, used to describe the variance of the gene counts as determined by:

$$Var\ K_{ij} = \mu_{ij} + \alpha_i \mu_{ij}^2$$

Each gene is taken to estimate gene-wise dispersion (maximum likelihood). In order to account for global variability in the data, the measure of the dispersion of the data counts was determined against the average expression for each experimental cohort. The dispersion parameter $\alpha_i$ follows a log-normal prior distribution that depends on the gene's mean normalized read count.

An empirical Bayes approach yields an estimate of how close the dispersions fit and the corresponding degrees of freedom. An ordinary GLM is performed to obtain maximum-likelihood estimates (MLEs) for the LFCs and then fit a zero-centered normal distribution to the observed distribution of MLEs over all genes. The data counts can be transformed using the regularized logarithmic transformation (*rlog*) by fitting each gene with a GLM. When we want to estimate the dispersion for a specific gene, we begin by fitting a negative binomial GLM to the count data for that gene. This initial fit uses a simple, approximate method called the method of moments to estimate the dispersion, which is based on the relationship between the average count and the variability within different experimental groups. This model does not include a log-fold change (LFC) prior in its design. The initial GLM is necessary to obtain an initial set of fitted values. We then maximize the Cox–Reid adjusted likelihood of the dispersion, conditioned on the fitted values from the initial fit, to obtain the gene-wise estimate, i.e., the initial GLM is necessary to obtain an initial set of fitted values, $\widehat{\mu_{ij}^{0}}$. We then maximize the Cox–Reid adjusted likelihood of the dispersion, conditioned on the fitted values $\widehat{\mu_{ij}^{0}}$ from the initial fit, to obtain the gene-wise estimate $\alpha_i^{gw}$, i.e.,

$$\alpha_i^{gw} = arg\ arg\ max_{\alpha}\ \iota_{CR}\left(\alpha; \overrightarrow{\mu_{\iota.}^{0}}, \overrightarrow{K_{\iota.}}\right)$$

With

$$\iota_{CR}\left(\alpha; \overrightarrow{\mu_{\iota.}^{0}}, \overrightarrow{K_{\iota.}}\right) = \iota(\alpha) - \frac{1}{2} log\ log\ (det\ det\ (X^{t}WX)\ )$$

$$\iota(\alpha) = \sum_{j} \quad log\ log\ f_{NB}\left(K_j;\ \mu_j, \alpha\right),$$

where $f_{\mathrm{NB}}(k;\mu,\alpha)$ is the probability mass function of the negative binomial distribution with mean $\mu$ and dispersion $\alpha$, and the second term provides the Cox–Reid bias adjustment.

The *rlog* transformation accounts for variation in sequencing depth across samples. The maximum a posteriori (MAP) as the final estimate of dispersion is plotted for the experimental groups (control versus pre-ALS participant cohort). The genes found in the MAP to be significant in expression were determined to be less than a false discovery rate of 10%. The standard outlier after the GLM fit is determined by the Cook's distance, defined as the scaled distance that the coefficient vector of the GLM would move if the gene in a sample were removed.

$$D_{ij} = \frac{R_{ij}^2}{\tau p} \frac{h_{jj}}{(1 - h_{jj})^2},$$

where $R_{ij}$ is the Pearson residual of sample $j$, $\tau$ is an overdispersion parameter (in the negative binomial GLM, $\tau$ is set to 1), $p$ is the number of parameters including the intercept, and $h_{jj}$ is the $j$th diagonal element of the hat matrix $H$:

$$H = W^{\frac{1}{2}}X(X^{t}WX)^{-1}X^{t}W^{\frac{1}{2}}.$$

Pearson residuals $R_{ij}$ are calculated as

$$R_{ij} = \frac{(K_{ij} - \mu_{ij})}{\sqrt{V(\mu_{ij})}}$$

where $\mu_{ij}$ is estimated by the negative binomial GLM without the LFC prior, and using the variance function $V(\mu)=\mu+\alpha\mu^2$.

The model coefficient from the GLM fit for each gene is compared to a level of zero. A Wald test is used to determine the significance of the LFC, the shrunken LFC estimate is divided by the error to yield a z-statistic and compared to a standard Gaussian distribution. The p values thus obtained are adjusted for multiple testing using the standard Benjamini and Hochberg statistical tests.

#### SM05. Weighted gene co-expression network analysis

A major caveat for differential analysis like DESeq2 is that it treats each gene as a separate entity and assumes independence of genes. However, in order to gain insight into complex biological mechanisms of regulation and co-expression, the genes were treated not as individual entities but as part of a larger group, co-expressed across the two experimental cohorts (control versus pre-ALS participant cohort). In order to visualize the changes in genes as modules of a network, we used the weighted gene co-expression network analysis (WGCNA) pipeline to analyze the gene groups in modules (https://cran.r-project.org/web/packages/WGCNA/index.html) (*10*). The top 30% of the most variable genes in the entire dataset were selected across the experiments using a standard variance calculation across the data points for each gene. For a ($n \times n$) count matrix, an adjacency matrix is measured as a weighted Pearson correlation:

$$s_{ij} = \frac{1 + corr(gene_i,\ gene_j)}{2}$$

Relying on the adjacency matrix, clustering is based on proximity of pairs of genes. WGCNA uses a topological overlap measure (TOM) which replaces the pairwise elements $s_{ij}$ by:

$$TOM_{ij} = \frac{\sum_{k\neq i,j} \quad s_{ik}s_{kj} + \alpha ij}{min(\sum_{k\neq i,j} \quad s_{ik,} \sum_{k\neq i,j} \quad s_{kj}) + 1 - s_{ij}}$$

Similarity of two genes *i,j* is determined by the strength of their interactions with the same neighbor genes. To enhance the strength of the correlations, a power transformation is applied:

$$a_{ij} = s_{ij}^{\beta}, \beta > 1$$

The power of the system networks is characterized by a power-law distribution, called scale free, with the adjacency equally distributed on [0,1]. A soft-power threshold of 0.12 is applied and the connectivity of the network, thus obtained, is defined as the total mean of the final adjacency matrix (TOM):

$$CC(A) = \frac{1}{n(n-1)} \sum_{i} \sum_{j \neq i} TOM_{ij}$$

Modules of genes are constructed by applying hierarchical clustering based on the TOM adjacency matrix. Each module is computed by taking the first principal component (PC) and designated as an eigen-gene. The whole network connectivity distribution is shown as a network heatmap, with the branches in the hierarchical clustering dendrograms corresponding to the modules/clusters of genes. We used the following parameters for our analysis:
Minimum module size (minMod)= 25;
Power of scale free network (ds) =3;
Dynamic cut height= 0.9999;
Soft threshold =0.12.
The WGCNA clustering yielded a total of 8 clusters with the control and pre-ALS participant cohorts analyzed. These clusters of genes are the ones that show correlated expression patterns across the control and ALS cohorts. In order to categorize the clusters further, we selected the main trends of the gene expression variability. Using the mean of the cluster Z-scores obtained for each gene in the WGCNA, we selected the clusters that showed similar trends across the analyzed groups. These scores were tested for statistical significance using Student's t-test on the cluster z-scores across the conditions: control versus pre-ALS participant cohort. The responsive trends across all the clusters are summarized as a heatmap. In order to account for the degree of similarity between the components in the categories and cross-sectional similarities, an autocorrelation of the obtained clusters was performed. The number of genes present in each category (the size of the category) determines the strength of that category, and the relative frequency of the genes present in that category is represented as a pie-chart.

#### SM06. Assessment of key nodes within the network - identifying key clusters and gene ranking for functional relevance

To assess the importance of individual nodes within the network we employed a metric based on the degree centrality. Degree centrality quantifies the average number of connections per node (degree) in the network. A higher average degree node signifies a more frequent exchange of information or interactions between the elements. We then evaluated the impact of each node on the overall network connectivity using a systematic removal approach. Each node was removed from the network and the average degree was recalculated. The difference between the modified and the original degree reflects the influence of each node. Nodes with a larger negative impact (greater decrease in average degree upon removal) were considered more central and crucial for maintaining the network's integrity. These nodes are hypothesized to play a more prominent role in regulating the cellular process under investigation.

This metric was applied to a cluster-cluster interaction map to identify key clusters which play a central role in the observed biological process.

Additionally, the metric was also applied in the context of ranking individual genes within the gene-gene network. This ranking provides an ordered list of genes based on their relative importance within the observed network. Individual genes with negative degree changes were defined as central genes.

#### SM07. Network analysis and enriched pathway functional characterization of central genes

Starting from the list of ordered genes within a cluster, only central genes were selected for further analysis. These genes were submitted to g:Profiler (https://biit.cs.ut.ee/gprofiler/gost, (*11*)) for ordered enrichment analysis. The resulting enriched pathways and central genes associated with them were used to construct a new interaction graph in a similar fashion as described before. Three network centrality measures were then applied to this pathway map degree centrality (number of connections), betweenness centrality (bridging subclusters of nodes within the network) and eigenvector centrality (connectivity to the most important nodes). Pathways with high scores across all centralities were considered as highly active drivers of the disease. High degree and eigenvector centrality suggested highly "impactful regulators of the disease", while high eigenvector and betweenness centrality pointed towards "critical connectors" crucial for signal transmission. To assess role of each pathway, each centrality measure was first normalized in the [0-1] range, prior to being multiplied based on predefined rules:

$$disease\ driver \rightarrow score\ I = degree \cdot betweenness \cdot eigenvector$$

$$key\ regulator \rightarrow score\ II = degree \cdot eigenvector$$

$$critical\ connector \rightarrow score\ III = betweenness \cdot eigenvector$$

#### SM08. Metabolic gene-metabolite network construction and analysis

The list of metabolic genes (C7- "metabolic pathways" associated genes) was subject to an integrated multi-omics analysis. Gene-metabolite connections were identified using MetaboAnalyst (https://www.metaboanalyst.ca/ ) by uploading the gene list and performing analysis against the KEGG Homo sapiens pathway database (https://www.kegg.jp/kegg/ ). MetaboAnalyst's internal algorithms identified direct reactions and associations. The resulting gene-metabolite network was quantitatively analyzed, with the top 10 genes and metabolites exhibiting the highest connectivity subsequently identified. Furthermore, enrichment analysis of the metabolic genes against KEGG and REACTOME (https://reactome.org/) metabolic pathways was performed in MetaboAnalyst as well as the enrichment analysis of transcription factor binding motifs.

#### SM09. Gene ontology analysis

Using the 4 categorized pools of genes that change in a particular fashion across the experimental groups, we performed a gene ontology (GO) analysis. The genes are annotated using annotator sets dedicated to the genes. GO leads to a list of significant shared GO terms (or parents of GO terms) used to describe the set of genes, the background frequency, the sample frequency, expected p-value, an indication of over/underrepresentation for each term, and p-value. We used the publicly available protocol in Metascape (www.metascape.org/) (*12*). GO analysis was focused on the following: GO database for biological processes, GO database for cellular components, GO database for molecular components, KEGG pathways and REACTOME pathways The minimum overlap was kept at 3, the p value cutoff at 0.01 and the minimum enrichment was kept at 1.5. For the mapping of interactome networks, we used String-db (https://stringdb.org). The network type was set at full network, and network edges were defined by confidence of the highest threshold (90%). For specific compartments of enrichment and processes for representation, we selected the pathways from the GO term for cellular compartment, biological processes, KEGG and REACTOME pathway descriptors.

#### Supplementary Figures

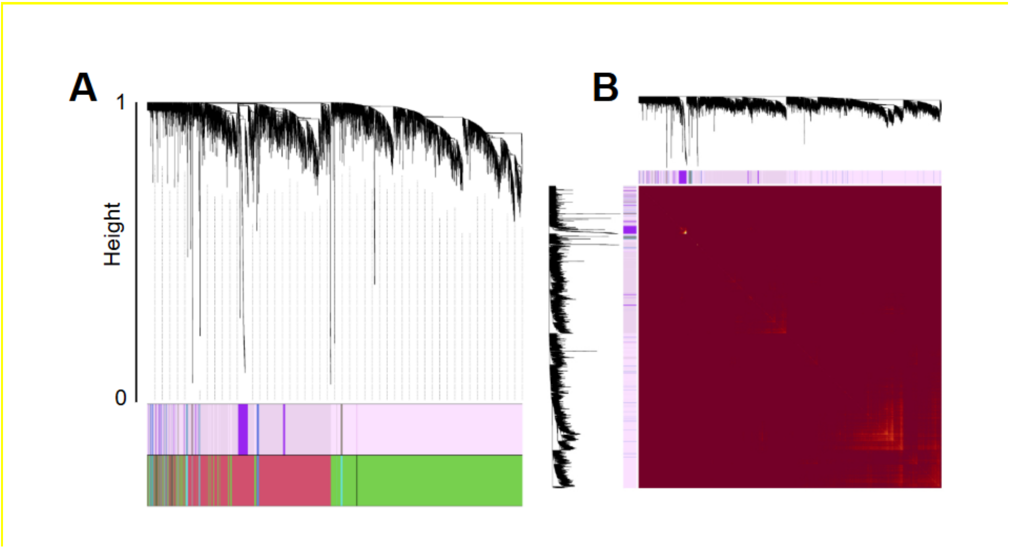


**Fig S1. Clustering of genes indicates key pathways segregated into two broad categories**
A. Dendrogram of the clusters obtained by WGCNA. (B) Topological overlap measure showing presence of key nodes in the clustered gene sets across control and PRE-ALS participants. Clusters with fewer than 50 genes (clusters 2 and 3) were excluded from further analysis.

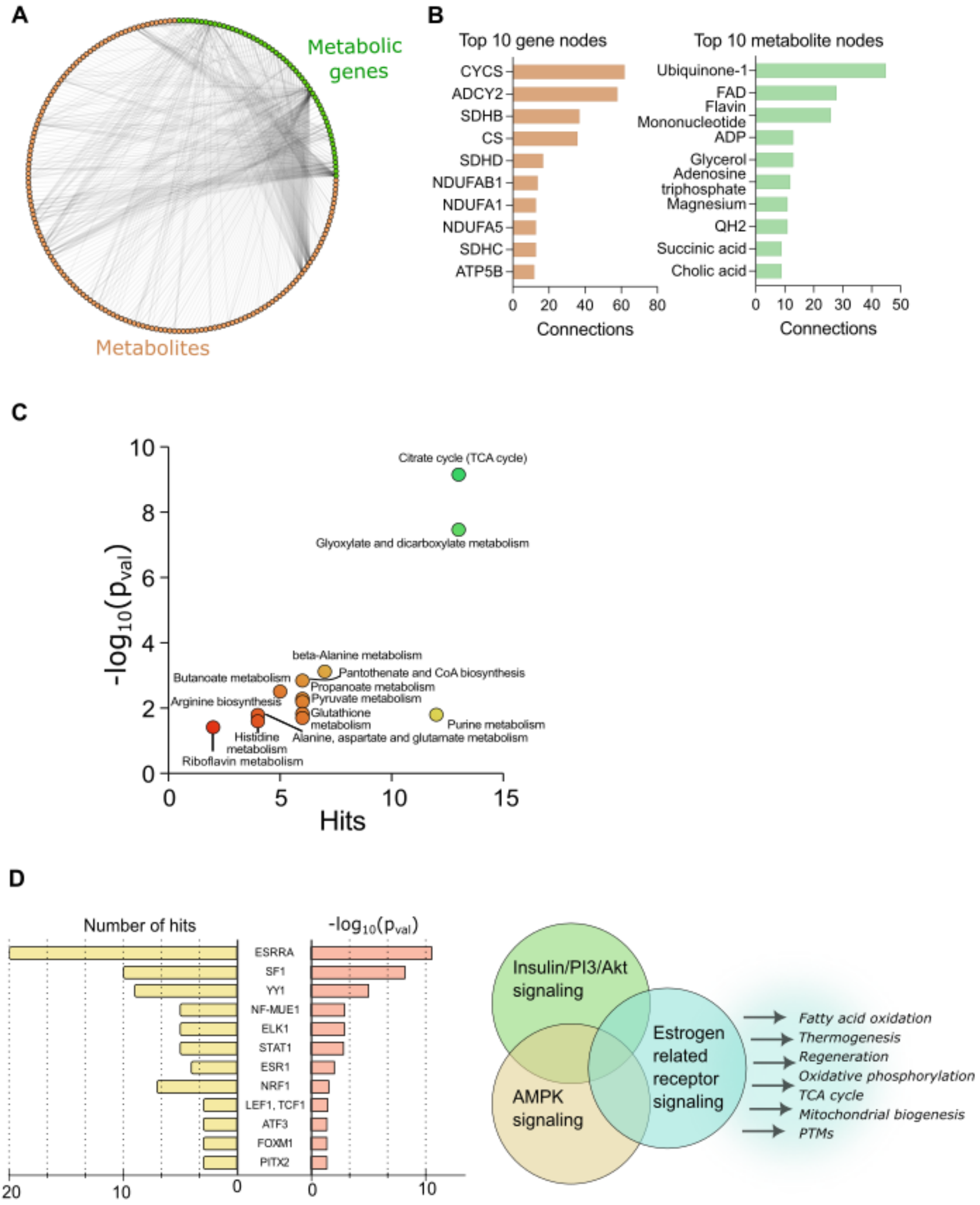


**Fig S2**. **Metabolic pathway depth in analysis (A)** Gene-metabolite network illustrating connections within the "metabolic pathway" as identified by MetaboAnalyst. **(B)** A focused view on the ten most highly connected genes and metabolites from the network in (A), indicating central players. **(C)** Results from KEGG metabolic enrichment analysis performed by MetaboAnalyst, where higher x- and y-axis values signify statistical significance of the enrichment and more hits (the number of genes identified within each enriched metabolic pathway), respectively. **(D)** Transcriptional motif enrichment analysis identifies ESRRA as a prominently enriched transcription factor. Literature review indicates AMPK and insulin signaling as upstream regulators of ESRRA (*13*), which in turn influences critical biological processes such as the TCA cycle, oxidative phosphorylation, post-translational modifications (PTMs), and thermogenesis (via UCP3).

Associated data:

- Datafile07_metabolic_enrichment.xlsx

A

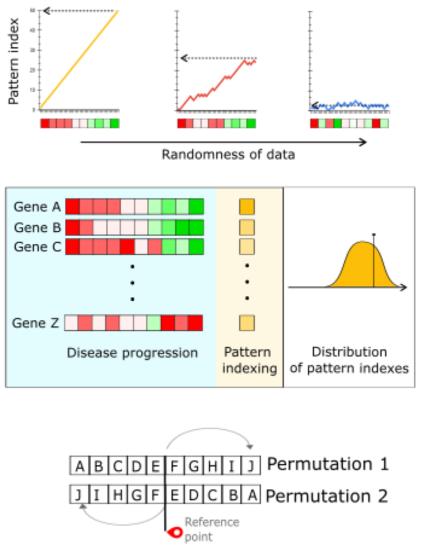


B

Distance from the reference point

| participant | 0 | 1 | 2 | 3 | 4 | 5 |
|---|---|---|---|---|---|---|
| A | 0 | 0 | 0 | 0.33 | 0.33 | 0.33 |
| B | 0.33 | 0 | 0 | 0.33 | 0.33 | 0 |
| C | 0 | 0.33 | 0.67 | 0 | 0 | 0 |
| D | 0.33 | 0.67 | 0 | 0 | 0 | 0 |
| E | 0.33 | 0.33 | 0 | 0 | 0 | 0.33 |
| F | 0 | 0.33 | 0.33 | 0 | 0.33 | 0 |
| G | 0 | 0.33 | 0.67 | 0 | 0 | 0 |
| H | 0 | 0.33 | 0.33 | 0.33 | 0 | 0 |
| I | 0 | 0 | 0 | 0.33 | 0.67 | 0 |
| J | 0 | 0 | 0.33 | 0 | 0.33 | 0.33 |

**Fig S3. Permutation of participant ordering**

Analysis of participant ordering permutations. A total of 100,000 participant order permutations were scanned (raw data in Datafile09_Tested_Permutations.txt). Three highly ordered gene expression patterns (permutations 1, 2, and 3) were identified for the selected key pathway genes. The probability of the distance from the array's center was calculated, accounting for the essential equivalence of reverse orders (e.g., ABC ≈ CBA). Probabilities and the three qualifying permutations are detailed in the xlsx table: (Datafile08_Ordered_Permutations.xlsx)

The pattern index effectively captures optimal participant stratification, with participants ordered along the x-axis based on the consistency of gene expression (either overexpression or downregulation) across the ordered list of expressed genesFor each randomly generated permutation, a pattern score was calculated for every gene by summing the changes in gene expression across consecutive samples (+1 for an increase, -1 for a decrease). The absolute value of this sum was used as the final score, where higher values indicated a more consistent monotonic trend. Permutations with a high-order probability (defined as the fraction of genes with a pattern score of 8 or greater) exceeding 0 were selected.

(A) Participants are ordered along the x-axis based on their likelihood of progressing to clinically-manifest ALS. Pattern index quantifies the consistency of monotonic gene expression changes across an ordered list of genes, with a high score indicating a strong pattern.

(B) To identify the optimal patient stratification for gene-conversion, we evaluated 100,000 participant permutations. A pattern score was calculated for all genes, and only permutations with a consistently high score (>8) were selected. Three optimal patterns were identified. Distances were calculated for each participant to determine the most probable position, revealing a progression from healthy to disease. This stratification aligned with our model, with individuals A and J representing the extremes of the disease spectrum. Remaining participants also followed the order predicted by our model.

*Associated data:*

- Datafile08_Ordered_Permutations.xlsx
- Datafile09_Tested_Permutations.txt
- Table S5

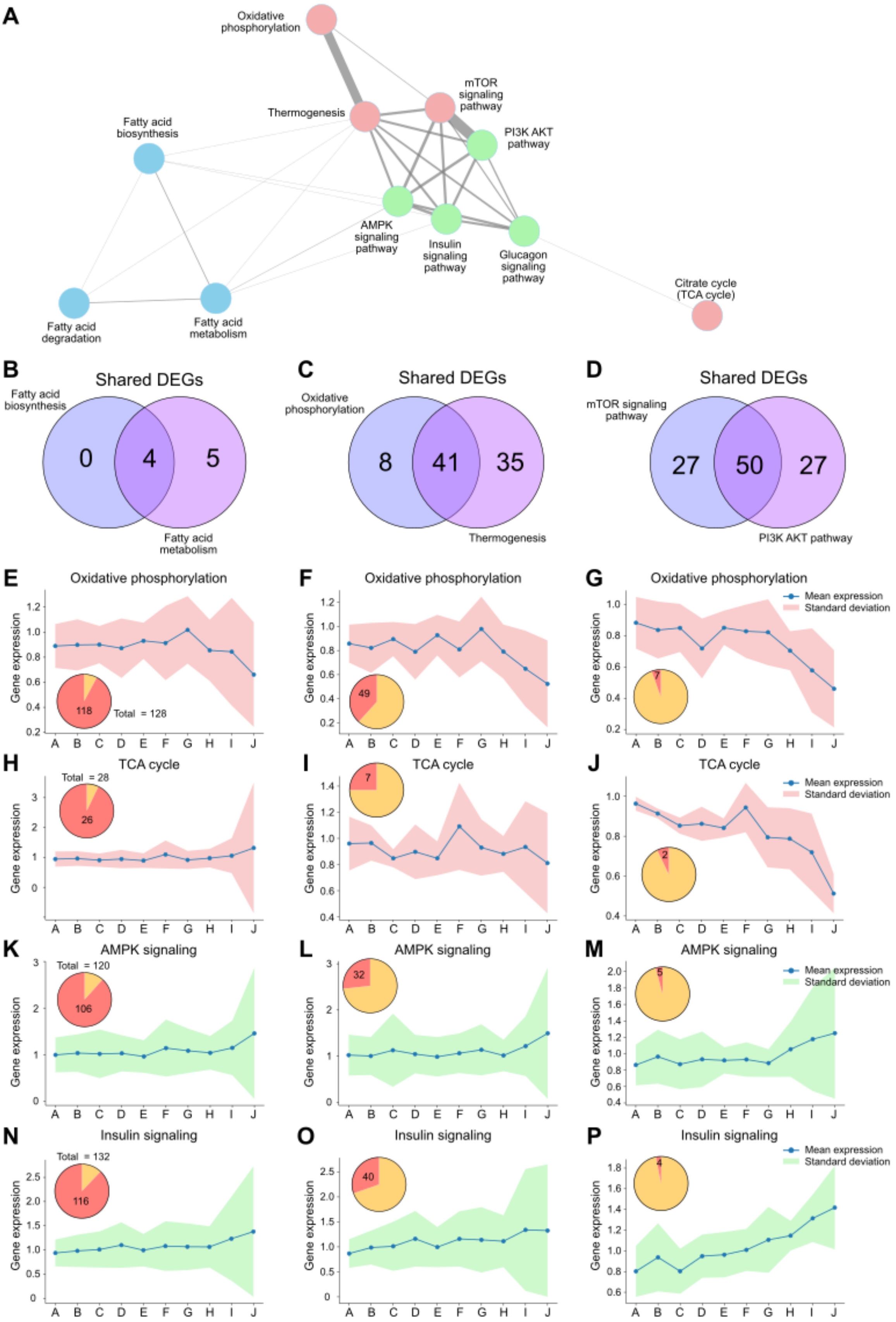

A
Oxidative phosphorylation
Thermogenesis
mTOR signaling pathway
PI3K AKT pathway
Fatty acid biosynthesis
AMPK signaling pathway
Insulin signaling pathway
Glucagon signaling pathway
Citrate cycle (TCA cycle)
Fatty acid degradation
Fatty acid metabolism
B
Shared DEGs
Fatty acid biosynthesis
0
4
5
Fatty acid metabolism
C
Shared DEGs
Oxidative phosphorylation
8
41
35
Thermogenesis
D
Shared DEGs
mTOR signaling pathway
27
50
27
PI3K AKT pathway
E
Oxidative phosphorylation
118
Total = 128
F
Oxidative phosphorylation
49
G
Oxidative phosphorylation
7
Mean expression
Standard deviation
H
TCA cycle
Total = 28
26
I
TCA cycle
7
J
TCA cycle
2
Mean expression
Standard deviation
K
AMPK signaling
Total = 120
106
L
AMPK signaling
32
M
AMPK signaling
5
Mean expression
Standard deviation
N
Insulin signaling
Total = 132
116
O
Insulin signaling
40
P
Insulin signaling
4
Mean expression
Standard deviation
Gene expression
A B C D E F G H I J

**Fig S4. Interplay and overlap of dysregulated pathways in PRE-ALS participants.**

To investigate pathway behavior, we employed an 'accumulator' metric, which quantifies the consistency of a gene's expression trend across an ordered list of participants by incrementing or decrementing its value by 0.1 for each subsequent participant with higher or lower expression, respectively. For our analysis, we selected genes based on their absolute accumulator values of 0, 0.2, and 0.4.

(A) We constructed a weighted network graph to visualize pathway interplay with nodes representing pathways and edges weighted by the number of shared genes meeting the accumulator criteria (acc ≥ 0.2). Node colors denote functional roles: red (energy production/anabolism/heat generation), green (key metabolic regulatory and signaling pathways), blue (lipid metabolism). Edge thickness reflects the number of shared genes exhibiting consistent and pronounced expression trends across participants, defined by an (|acc|) ≥0.2, a metric that quantifies the consistency and magnitude of directional gene expression changes across participants. (B-D) Venn diagram highlighting the significant gene overlap between fatty acid metabolism and fatty acid biosynthesis pathways (|acc| ≥ 0.2) (B), oxidative phosphorylation and thermogenesis (C) and mTOR and PI3-AKT pathways (|acc| ≥ 0.2) (D). (E-P) Fold change expression patterns across participants for oxidative phosphorylation, TCA signaling, AMPK and insulin signaling pathways (displayed row-wise). Genes included in each panel were selected based on their |acc| value: ~0 (E, H, K, N), 0.2 (F, I, L, O), and 0.4 (G, J, M, P), representing increasingly consistent expression trends. The pie charts show the proportion of genes that meet the accumulator value criteria (red) versus the remaining genes (yellow) within each pathway. Oxidative phosphorylation consistently showed downregulation across all accumulator criteria, from the most inclusive (acc ≈ 0) to the most stringent (acc = 0.4) gene sets (Fig. E-G). Pathways, including the TCA cycle, AMPK signaling, and insulin signaling, exhibited clear expression trends only under the most stringent criterion (acc = 0.4) (Fig. J, M, P). This trend weakens as less consistent genes are included (Fig. H, K, N), potentially due to shared pathway components or compensatory mechanisms.

### Supplementary Tables

| | Asymptomatic ALS gene carriers | | Control individuals (n= 10) | Statistics[&] |
|---|---|---|---|---|
| | *Baseline (n=10)* | *18 months (n=9)[#]* | | |
| **Age at inclusion (y)** | 48,4 +/- 11,4 [31-67] | | 47,9 +/- 10,6 [30-60] | NS |
| **Gender (female/male, n)** | 7/3 | | 7/3 | NS |
| **CMAP deltoid muscle (mV)** | 10,90 +/- 0,86 [9,67-12,48] | 11,97 +/- 1,94 [9,11-14,99] | 11,47 +/- 2,64 [7,53-16,26] | NS |
| **MUNIX deltoid muscle** | 243,80 +/- 28,37 [188-279] | 271,56 +/- 54,94 [152-355] | 251,10 +/- 78,45 [157-410] | NS |
| **CMAP sum-score (mV)*** | 44,99 +/- 2,79 [38,43-48,30] | 46,08 +/- 3,85 [40,51-51,74] | 45,33 +/- 6,35 [36,66-53,20] | NS |
| **MUNIX sum-score*** | 769,50 +/- 110,99 [586-945] | 827,67 +/- 122,66 [537-998] | 836,80 +/- 155,17 [599-1094] | NS |

**Table S1. Characteristics of PRE-ALS participants and control individuals for neurophysiological studies**

All PRE-ALS participants were free of symptoms suggestive of amyotrophic lateral sclerosis (ALS) or cognitive impairment. Neurological examination by an experienced neurologist (MDMA, FS, or GB) revealed no signs of motor neuron disease. Muscle strength was normal in all participants, with a total Medical Research Council (MRC) score of 130/130. Mini-Mental State Examination (MMSE) scores ranged from 28 to 30/30.

Control data were obtained from healthy subjects previously recorded in the Neurophysiology Laboratory of the Pitié-Salpêtrière Hospital. Data are presented as mean +/- SD [range] unless otherwise specified.

[#] At 18 months, electrophysiological recording could not be performed in one asymptomatic ALS gene carrier

[&] The Mann-Whitney test was used to compare electrophysiological results between asymptomatic ALS gene mutation carriers at baseline and control individuals. The Wilcoxon signed rank test was used to compare electrophysiological results in asymptomatic ALS gene mutation carriers between baseline and the 18-month follow-up visit.

* CMAP sum-score and MUNIX sum-score were calculated by summing the values of the 5 muscles tested.

CMAP: compound muscle action potential, MUNIX: Motor Unit Number Index

| Control individuals § | Age (years) | Sex (M/F) | Muscle |
|---|---|---|---|
| 1 | 44 | F | Deltoid |
| 2 | 46 | F | Deltoid |
| 3 | 45 | F | Deltoid |
| 4 | 45 | F | Deltoid |
| 5 | 45 | F | Deltoid |
| 6 | 53 | F | Deltoid |
| 7 | 60 | F | Deltoid |
| 8 | 62 | M | Deltoid |
| 9 | 58 | F | Deltoid |
| 10 | 56 | M | Deltoid |

**Table S2: Characteristics of control individuals for RNA seq analysis**

§ All controls underwent a diagnostic muscle biopsy for non-specific symptoms (pain and/or fatigue) and were considered free of any neuromuscular disorder after careful review of morphological and histochemical examinations.

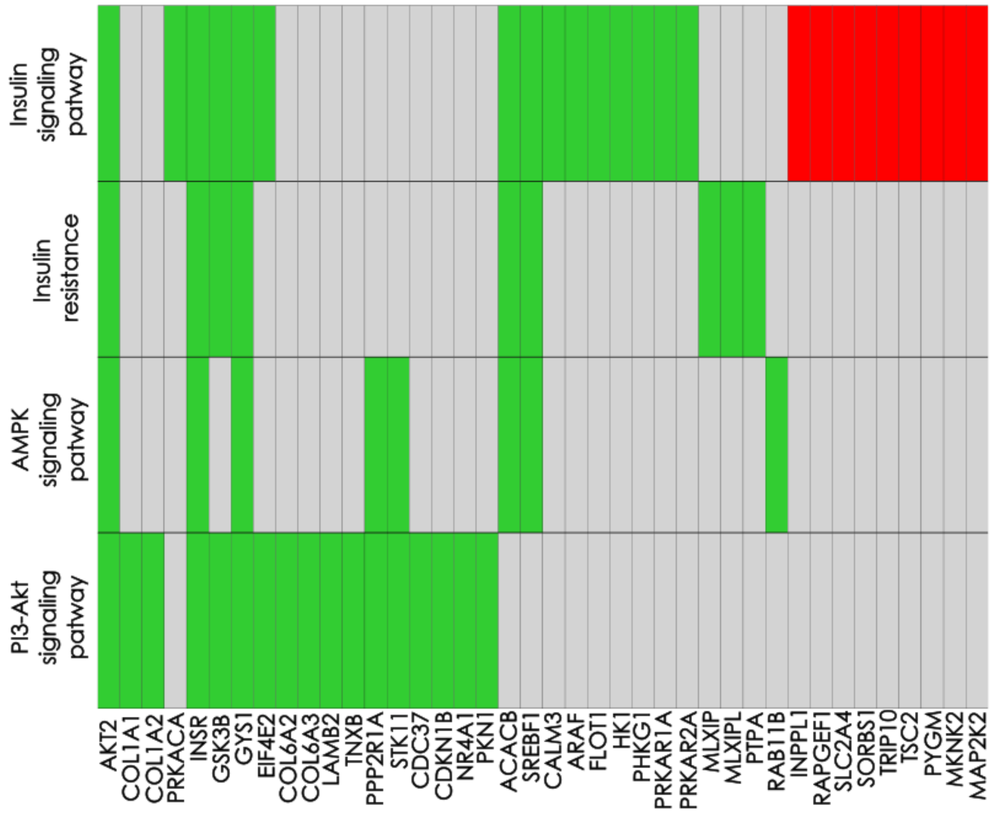


**Table S3. Genes belonging to selected C7 and C8 pathways (PI3-Akt, AMPK and insulin signaling and insulin resistance pathway).**

Green indicates upregulation, red indicates downregulation, and grey indicates absence from the pathway, as determined by DESeq2 analysis.

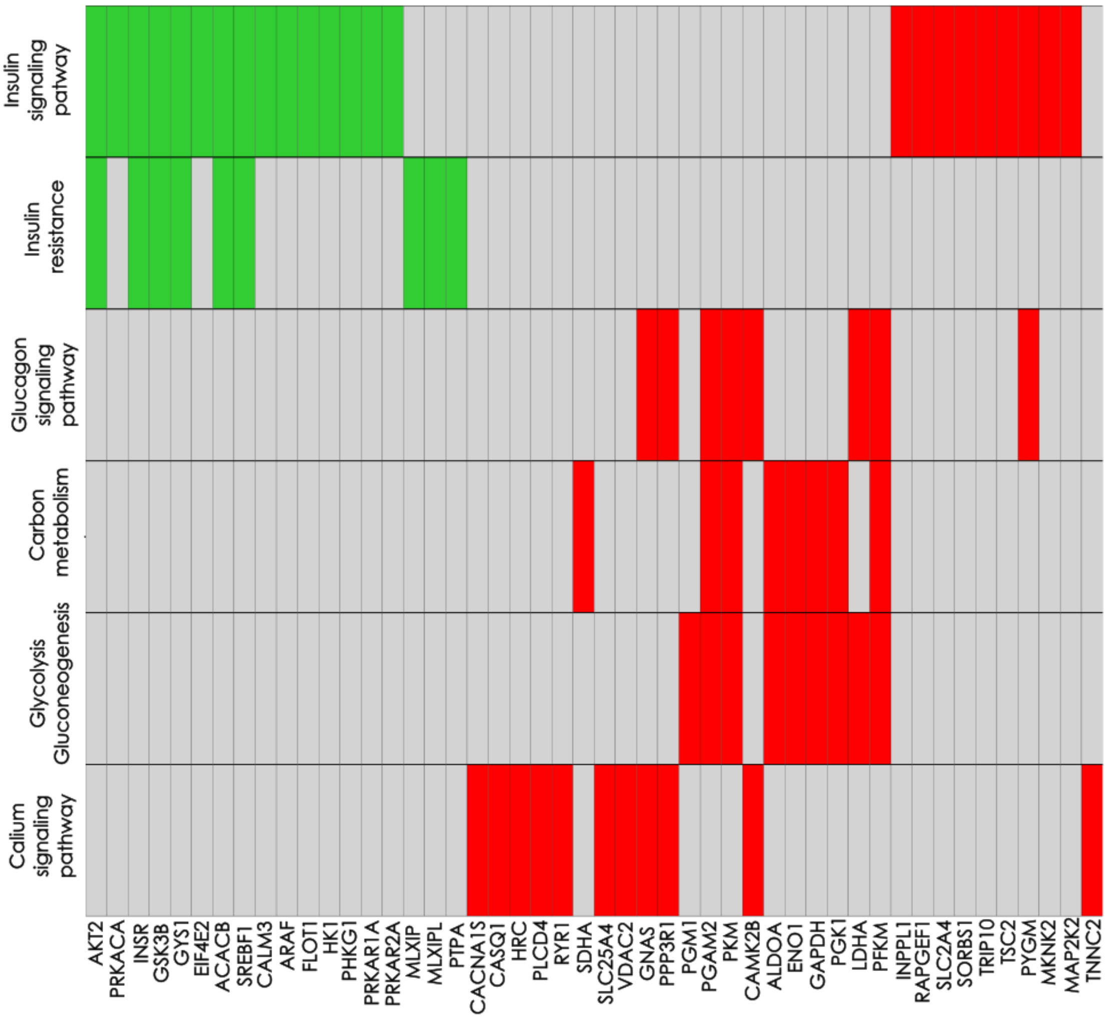


**Table S4**. **Genes belonging to selected C7 and C8 pathways (calcium, glucagon and insulin signaling pathways, carbon metabolism, insulin resistance and glycolysis/gluconeogenesis pathway).**

Green indicates upregulation, red indicates downregulation, and grey indicates absence from the pathway, as determined by DESeq2 analysis.

| position | 1 | 2 | 3 | 4 | 5 | 6 | 7 | 8 | 9 | 10 |
|---|---|---|---|---|---|---|---|---|---|---|
| permutation 1 | a | b | c | d | e | f | g | h | i | j |
| permutation 2 | j | i | g | c | d | e | h | f | b | a |
| permutation 3 | f | a | c | h | b | d | j | g | i | e |
| **abs value ( position-5)** | **4** | **3** | **2** | **1** | **0** | **1** | **2** | **3** | **4** | **5** |

**Table S5:** Three highly ordered gene expression patterns (permutations 1, 2, and 3)

### Supplementary information for additional datafiles

##### SI1. Normalized transcript counts, fold change calculation, and cohort heterogeneity

Differential expression analysis results (Datafile01_diffexpr-results_counts.xlsx) utilized normalized transcript counts. The fold change values (Datafile02_fold_change.xlsx) presented throughout the main manuscript [Figure 6I-M, figure 7 and figure S4] reflect the ratio of the normalized expression counts in PRE-ALS participants relative to the mean normalized expression counts of the control group.

As discussed in detail in the main text, it is critical to consider the inherent heterogeneity of the PRE-ALS participant group. This cohort is special because it includes individuals who carry the relevant mutation but may remain clinically healthy, while some are progressing toward the disease state—and those who are progressing are not necessarily at the same stage. In contrast, the control group represents a more homogeneous population. This acknowledged difference in group structure is accounted for in the interpretation of all differential expression analyses.

The correlation analysis between upstream and downstream genes within the main text [figure 6] was performed using these normalized counts [Datafile01_diffexpr-results_counts.xlsx ] mentioned here]. For example, the correlation analysis involving autophagy (PPARGC1A, SREBF1, TSC2, AKT2) can be found in [Datafile04_PGC1alpha_deactivation - PGC1alpha.xlsx].

Associated data:

Datafile01_diffexpr-results_counts.xlsx

Datafile02_fold_change.xlsx

Datafile04_PGC1alpha_deactivation - PGC1alpha.xlsx

##### SI2. Network analysis and pathway enrichment of key clusters and their central genes

The importance of a node (gene) within its cluster was determined by calculating its degree change in the network. Based on the genes belonging to identified clusters, a cluster interaction map was constructed, and the degree change was calculated for every cluster.

For the two most significant clusters, 7 and 8, central genes were pinpointed by selecting those exhibiting a negative degree change (degree change <0). Separately for both up- and downregulated central genes, pathway enrichment analysis was performed using g:Profiler. All disease-related pathways were manually excluded from the final enrichment results to focus on core biological functions

For a full explanation of how the degree change metric was calculated, refer to the supplementary methods section titled "Assessment of key nodes within the network - identifying key clusters and gene ranking for functional relevance."

Associated data:

-Datafile05_preALS_supplementary_figure 3.xlsx

**SI3. Analysis of key central genes, centrality metrics of C7 and C8**

Central genes identified in clusters 7 and 8 were prioritized by ordering them from the most central to the least central. Gene importance within enriched pathways was assessed by calculating three network centrality metrics based on the pathway interactome: degree, betweenness, and eigenvector centrality.

Centrality values were first normalized, and then their coefficients were calculated. The statistical significance of each pathway was determined using the Kolmogorov-Smirnov (KS) statistic to calculate a p-value comparing gene expression counts between control and pre-treatment participants (control versus PRE-ALS counts). Only pathways meeting the significance threshold of $p<0.05$ were carried forward for further analysis. Details on the calculation of the centrality coefficients can be found in the supplementary methods section titled "Network analysis and enriched pathway functional characterization of central genes."

Associated data:

-Datafile06_supplementary_figure 4,5.xlsx

**SI4. Overlap analysis of dysregulated pathways in PRE-ALS participants**

Excel sheet 'Datafile03_Pathways and gene associations_supplementary figure 4.xlsx ' details the associations between pathways and individual genes, as visualized in Figure S4. The analysis focuses on the interplay and functional overlap of the identified dysregulated pathways in the PRE-ALS participant group for different Acc values.

Associated data:

Datafile03_Pathways and gene associations_supplementary figure 4.xlsx
